# The activation of mGluR4 rescues parallel fiber LTP, motor learning and social behavior in a mouse model of Fragile X Syndrome

**DOI:** 10.1101/2022.04.29.490046

**Authors:** Ricardo Martín, Alberto Samuel Suárez-Pinilla, Nuria García-Font, M. Luisa Laguna-Luque, Juan C. López-Ramos, María Jesús Oset-Gasque, Agnes Gruart, José M. Delgado-García, Magdalena Torres, José Sánchez-Prieto

## Abstract

Fragile X patients and mice lacking the Fragile X Mental Retardation Protein (FMRP) suffer from multiple behavioral alterations, including deficits in motor learning. We found that enhanced synaptic vesicle (SV) docking in cerebellar parallel fiber to Purkinje cell *Fmr1*KO synapses was associated with enhanced asynchronous release, which not only occludes further potentiation, but it also compromises presynaptic parallel fiber long-term potentiation (PF-LTP). A reduction in extracellular Ca^2+^ restored the readily releasable pool (RRP) size, rescuing β adrenergic receptor-mediated potentiation and parallel fiber LTP. Interestingly, VU 0155041, a selective positive allosteric modulator of mGluR4, also restored both the RRP size and parallel fiber LTP. Moreover, when injected into *Fmr1*KO mice, VU 0155041 improved motor learning in skilled reaching, classical eyeblink conditioning and vestibuloocular reflex (VOR) tests, as well as improving the social behavior of these mice. Thus, pharmacological activation of mGluRs may offer therapeutic relief for motor learning and social deficits in Fragile X Syndrome.

## 1. INTRODUCTION

Fragile X syndrome (FXS), the most common inherited intellectual disability, is associated with multiple behavioral alterations, such as cognitive deficits, hyperactivity, anxiety and impaired social interactions (Hagerman et al., 2009; MacLeod et al., 2010). FXS patients also experience deficits in the acquisition of motor skills, affecting fine and gross motor activities. (Will et al., 2018). FXS is caused by the silencing of the *Fmr1* gene, which encodes the fragile mental retardation protein (FMRP). FMRP is an RNA-binding protein that negatively regulates protein synthesis (Bassell and Warren, 2008) and in its absence, the expression of many postsynaptic proteins is altered, affecting long-term forms of postsynaptic plasticity (Huber et al., 2002). FMRP can be found in axons and presynaptic nerve terminals (Akins et al., 2012; Christie et al., 2009). Indeed, proteomic studies on a mouse model of FXS, *Fmr1*KO mice, revealed a presynaptic phenotype (Klemmer et al., 2011), with altered expression of presynaptic proteins involved in excitability, Ca^2+^ homeostasis and neurotransmitter release (Christie et al., 2009; Darnell et al., 2011, 2001; Liao et al., 2008). Electron microscopy (EM) studies of *Fmr1*KO synapses identified an increase in the number of docked synaptic vesicles (SVs) (Deng et al., 2011a, 2013; García-Font et al., 2019) that was driven by a series of factors. FMRP interacts with the Ca^2+^ activated K^+^ channels that control the duration of action potentials (APs) and thus, the loss of FMRP leads to AP broadening and an ensuing increase in Ca^2+^ influx (Deng et al., 2013). In addition to a higher Ca^2+^ influx, *Fmr1*KO cells also accumulate more diacylglycerol (DAG) due to the loss of the diacylglycerol kinase (Tabet el al., 2016). Both Ca^2+^ and DAG can enhance SV docking and priming through the activation of the Munc13 proteins that bind to these messengers (Rhee et al., 2002; Shin et al., 2010), and that may also bind to calmodulin (CaM) (Junge et al., 2004). Changes in SV docking could affect neurotransmitter release, as the number of docked vesicles is correlated with the size of the readily releasable pool (RRP) of SVs (Rosenmund and Stevens, 1996; Schikorski and Stevens, 2001). Specifically, the enhanced SV docking at *Fmr1*KO synapses may occlude presynaptic forms of synaptic plasticity, such as Long Term Potentiation (LTP) at cerebellar parallel fibers (PFs), which is expressed through an increase in neurotransmitter release.

LTP at PF to Purkinje cell (PF-PC) synapses depends on enhanced neurotransmitter release, which is mediated by a Ca^2+^-induced increase in presynaptic cAMP (Salin et al., 1996; Storm et al., 1998) and the active zone (AZ) protein RIM1α (Castillo et al., 2002). We recently found that PF-PC LTP requires a β-adrenergic receptor (β-AR) dependent increase in SV docking and an increase in the RRP size (Martín et al., 2020), events that contribute to enhanced neurotransmitter release (Martín et al., 2020; Ferrero et al., 2013). However, the potentiation of release by β-ARs is impaired in *Fmr1*KO synaptosomes (García-Font et al., 2019). Plasticity at PF-PC synapses is involved in cerebellar motor learning (Ito, 2002; Schonewille et al., 2010; Gutierrez-Castellanos et al., 2017) and this may be related to the motor learning deficits evident in *Fmr1*KO mice (Padmashri et al., 2013). Thus, we hypothesized that rescuing the eventual loss of PF-PC LTP in *Fmr1*KO mice could help ameliorate their motor learning deficits (Padmashri et al., 2013).

Here we found that PF-PC LTP is lost in *Fmr1*KO synapses because they have more docked SVs in the basal state and a larger RRP size than wild type (WT) synapses, such that β-AR mediated potentiation is occluded. Lowering extracellular Ca^2+^ ([Ca^2+^]_e_) to 1 mM restored SV docking, the RRP size and PF-PC LTP. These ameliorating effects were also produced by the selective positive allosteric modulator (PAM) of mGluR4, VU 015504, which reduces Ca^2+^ influx at nerve terminals. Interestingly, the motor learning of *Fmr1*KO mice that received VU 0155041 improved, as did their social interactions. Thus, pharmacological activation of mGluR4 may offer therapeutic relief from the motor and behavioral deficits in FXS.

## 2. RESULTS

### 2.1. The lack of FMRP increases aEPSC frequency and occludes isoproterenol induced potentiation despite normal β-AR expression and cAMP generation

There is enhanced SV docking at synapses in *Fmr1*KO mice (Deng et al., 2011a) and thus, we asked whether this results in an increase in spontaneous neurotransmitter release. However, as it is not possible to distinguish PF from climbing fiber miniature excitatory postsynaptic currents (mEPSCs), we measured asynchronous release in the presence of Sr^2+^. Under these conditions, asynchronous release represents single release events, and it is associated with PF stimulation (Carey and Regehr, 2009). In WT slices, isoproterenol increased synchronous release (*P*<0.0001, Fig. 1*A,B*) and the frequency (*P*<0.0001, Fig. 1*C,D,E*) but not the amplitude (*P*=0.7443, Fig. 1*C,F,G*) of asynchronous release. However, isoproterenol failed to enhance sEPSCs at *Fmr1KO* slices (*P*=0.4958, Fig. 1*H,I*), and as the frequency of asynchronous aEPSCs in *Fmr1KO* mice was higher than in WT slices (*P*=0.0001, Fig. 1*D,K*), this further occluded the potentiation by isoproterenol (*P*=0.8764, Fig. 1*J,K,L*).

We assessed whether the failure of isoproterenol to potentiate neurotransmitter release at *Fmr1*KO synapses was a consequence of changes in β-AR expression and/or activity, which was examined in a preparation of cerebellar nerve terminals (synaptosomes). The spontaneous release of glutamate from WT cerebellar synaptosomes increased in the presence of isoproterenol (*P*=0.0005, *Supplementary Material:* Fig. S1*A,C*). However, there was more spontaneous release in *Fmr1*KO synaptosomes under basal conditions (*P*=0.011, *Supplementary Material:* Fig. S1*C*) but this was not further potentiated by isoproterenol (*P*= 0.719, *Supplementary Material:* Fig. S1*B,C*). Thus, *Fmr1*KO cerebellar synaptosomes recapitulate the occlusion phenotype evident in *Fmr1*KO slices.

**Figure 1.**
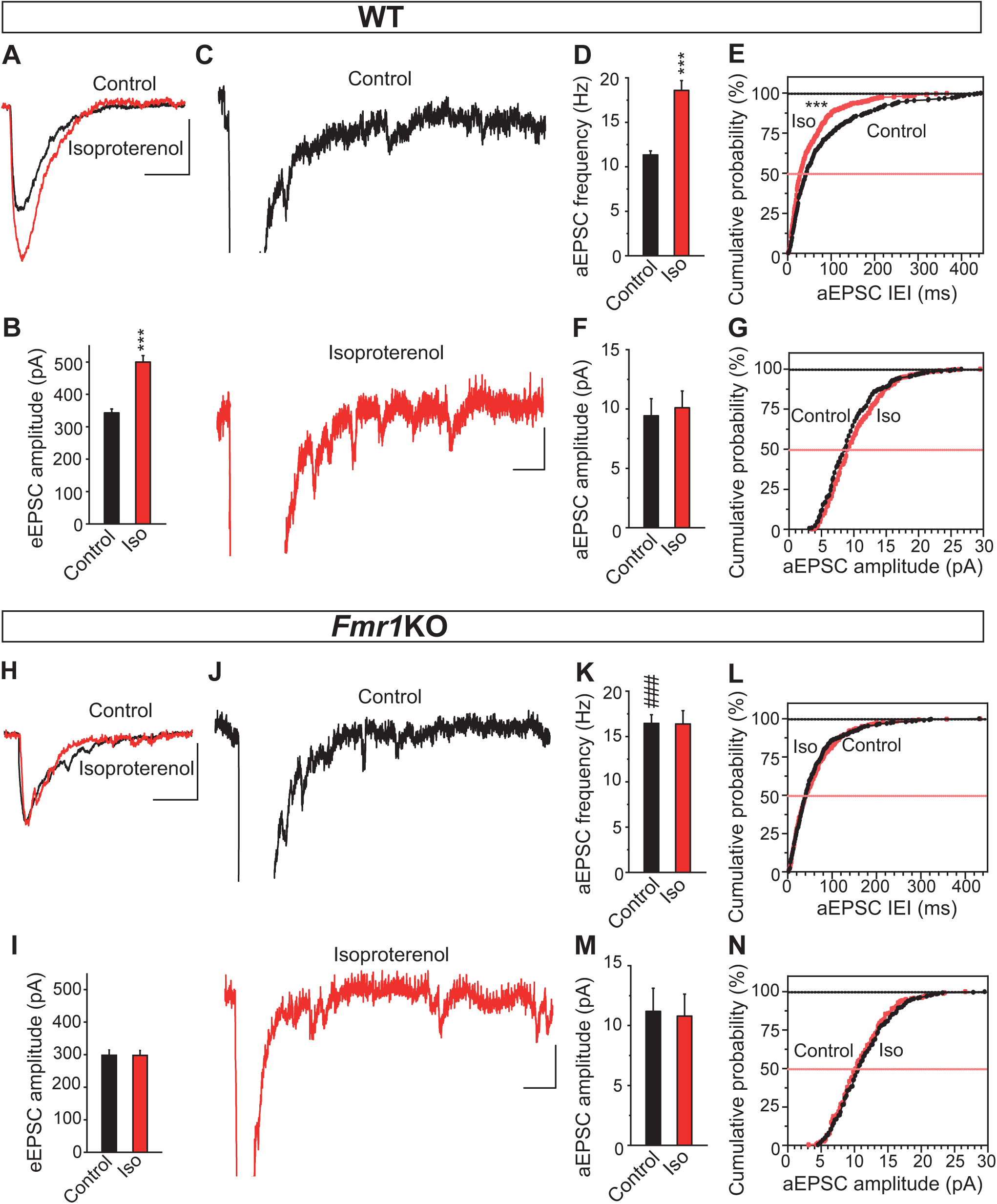
An increased aEPSC frequency and occlusion of isoproterenol induced potentiation at *Fmr1*KO synapses. (A, H) Isoproterenol (100 μM, 10 min) enhances the sEPSCs recorded in the presence of Sr^2+^ (2.5 mM) in WT (A) but not in *Fmr1*KO slices (H). (B, I) Quantification of the effects of isoproterenol on the sEPSC amplitude in WT (B) (n=9, *P*<0.0001) and in *Fmr1*KO slices (I) (n=6, *P*=0.4958). (C, J) Individual traces showing asynchronous release events in control (black) and isoproterenol exposed (red) WT (C) or *Fmr1*KO slices (J). (D, F, K, M) Quantification of the isoproterenol induced changes in aEPSC frequency in WT (D) (n=306 events/9 slices and, n=502 events/9 slices: *P*<0.0001) and *Fmr1*KO slices (K) (n=305 events/6 and n=312 events/6 slices: *P*=0.8764), as well as in amplitude in WT (F) (*P*=0.7443) and in *Fmr1*KO slices (M) (*P*=0.9463). (E, G, L, N) Cumulative probability plots of isoproterenol induced changes in aEPSC frequency (inter event interval, IEI) WT (E) (*P*=0.0003) and *Fmr1*KO slices (L), (*P*=0.7583), and amplitude in WT (G) (*P*=0.0885) and *Fmr1*KO (N) (*P*=0.4917). The data represent the mean ± SEM. Scale bars in (A, H) and (C, J) represent 100 pA and 10 ms, and 25 pA and 10 ms, respectively. Unpaired student’s t test in (B,D,F,I,K,M). Kolmogrov-Smirnov test in (E,G,L,N). \*\**P*<0.01, \*\*\**P*<0.001.

The absence of β-AR mediated potentiation of spontaneous release in *Fmr1*KO synaptosomes could be due to weaker expression of this receptor. However, β1-AR expression was similar in *Fmr1*KO and WT synaptosomes when assessed in western blots (*P*=0.5773, *Supplementary Material:* Fig. S2*A,B)*. We also assessed β-AR expressing synaptosomes by immunofluorescence using antibodies against β_1_-AR and synaptophysin as a marker of SVs. There were a similar number of *Fmr1*KO and WT cerebellar synaptosomes labeled for synaptophysin that also expressed β_1_-AR synaptosomes (*P*=0.2000, *Supplementary Material:* Fig. S3*A,B,C*). These data indicate that a change in β_1_-AR expression is not responsible for the failure of isoproterenol to potentiate spontaneous release in *Fmr1* KO synaptosomes. β-ARs activate Adenylyl Cyclase (AC) and generate cAMP, which activates downstream signals to potentiate spontaneous release (Ferrero et al., 2013). The lack of isoproterenol induced potentiation was not due to loss of receptor function as there was a similar β-AR mediated increase in cAMP in *Fmr1*KO synaptosomes as that in WT synaptosomes (*P*=0.848, *Supplementary Material:* Fig. S4*A*), reinforcing the idea that potentiation mediated by β-ARs is occluded at *Fmr1* KO synapses.

### 2.2. *Fmr1*KO synapses have more docked SVs than WT synapses and they are insensitive to isoproterenol

The number of docked vesicles at PF-PC WT synapses (those located within 10 nm of the AZ membrane) were increased by exposure to isoproterenol (*P*<0.001, Fig. 2*A,C,D*). Notably, there were more docked SVs at PF-PC *Fmr1*KO synapses in the basal state than in WT synapses (*P*<0.001) and isoproterenol failed to significantly increase this parameter (*P*>0.05, Fig. 2*E,G,H*). Together, the failure of β-ARs to potentiate neurotransmitter release seems to be due to these receptors failing to enhance the docking of SVs to the plasma membrane AZ, which is already enhanced at *Fmr1*KO PF-PC synapses.

**Figure 2.**
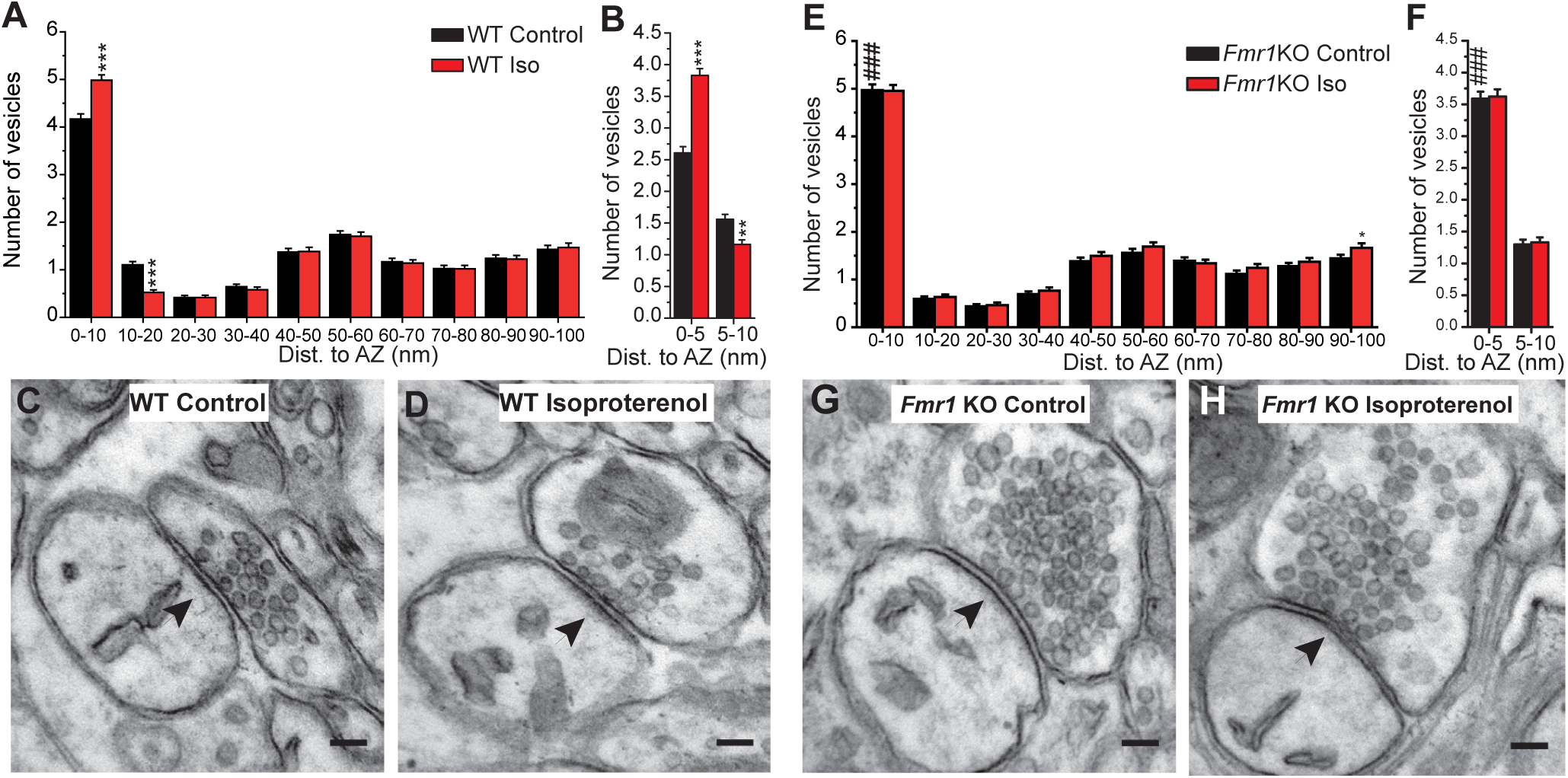
Enhanced SV docking of *Fmr1* KO synapses and lack of an effect of isoproterenol. Isoproterenol (100 μM, 10 min) increases SV docking in WT (A,C,D) but not in *Fmr1*KO (E,G,H) PF-PC synapses. (A) Isoproterenol changes SV distribution (0-10 nm) in WT (n=250/3 and 203/3 synapses/mice: *P*<0.001) but not in *Fmr1*KO slices (0-10 nm) (n=238/4 and 224/4 synapses/mice, *P*>0.05). Note the increase in SV docking in WT control as opposed to *Fmr1*KO control slices (^###^*P*<0.001). (C,D,G,H) Electron microscopy of PF-PC synapses. (B,F) The effect of isoproterenol on the SV distribution: WT (0-5 nm: *P*<0.001); WT (5-10 nm: *P*<0.01); *Fmr1*KO (0-5 nm: *P*>0.05) and *Fmr1*KO (5-10 nm: *P*>0.05). *Fmr1*KO control vs WT control (^###^*P*<0.001). The values represent the mean ± S.E.M. Scale bar in (C,D,G,H) 100 nm. Two-way ANOVA with Bonferroni in (A,B,E,F). \*\**P*<0.01, \*\*\**P*<0.001.

Loosely docked and primed SVs located 8 nm from the AZ membrane due to the partial zippering of SNARE complexes were distinguished from the tightly docked and fully primed SVs in which SNARE complex zippering has progressed much further, bringing the SVs closer to the AZ membrane (0-5 nm) (Fernández-Busnadiego et al., 2013; Imig et al., 2014; Taschenberger et al., 2016; Neher and Brose, 2018). There were more SVs within 0-5 mm of the AZ membrane in WT slices following exposure to isoproterenol (*P*<0.001) at the expense of SVs within 5-10 nm, which decreased in number (*P*<0.01, Fig. 2*B*). However, *Fmr1*KO synapses had more SVs within 5 nm than WT synapses (*P*<0.001, Fig. 2*F*) and isoproterenol failed to further increase this parameter (*P*>0.05). Isoproterenol did not change the SVs at a distance of 5-10 nm from the AZ membrane in *Fmr1*KO synapses (*P*>0.05, Fig. 2*F*). Thus, *Fmr1* KO synapses have more tightly docked and fully primed SVs than WT synapses, and the ability of β-ARs to further increase this parameter is occluded.

### 2.3. Cerebellar parallel fiber LTP is absent in PF-PC Fmr1 KO synapses

The enhanced SV docking and increased RRP size mediated by β-ARs is essential for PF-PC LTP, and indeed, β-AR antagonists prevent this presynaptic form of plasticity (Martín et al., 2020). Presynaptic LTP at PF-PC synapses depends on a Ca^2+^ mediated increase in presynaptic cAMP (Salin et al., 1996) and on the RIM1α protein at the AZ, a Rab-3 interacting molecule (Castillo et al, 2002). As presynaptic responses of β-ARs are occluded at *Fmr1*KO PF-PC synapses, we assessed whether PF-PC LTP was lost at these synapses. PF-PC LTP can be induced by stimulating PFs at 10 Hz for 10s (Castillo et al., 2002) and it is expressed as a long lasting increase in EPSC amplitude (*P*=0.0001, Fig. 3*A,B*). The presynaptic origin of this LTP was evident by the decreased paired-pulse ratio (PPR; *P*=0.0317, Fig. 3*C,D*), and the β-AR receptor antagonist propranolol prevented PF-PC LTP in WT slices (*P*=0.7125, Fig. 3*A,B*). However, stimulation of PFs at 10 Hz for 10 sec failed to induce LTP at PF-PC *Fmr1*KO synapses (*P*=0.8955, compared to the baseline: Fig. 3*A,B*).

**Figure 3.**
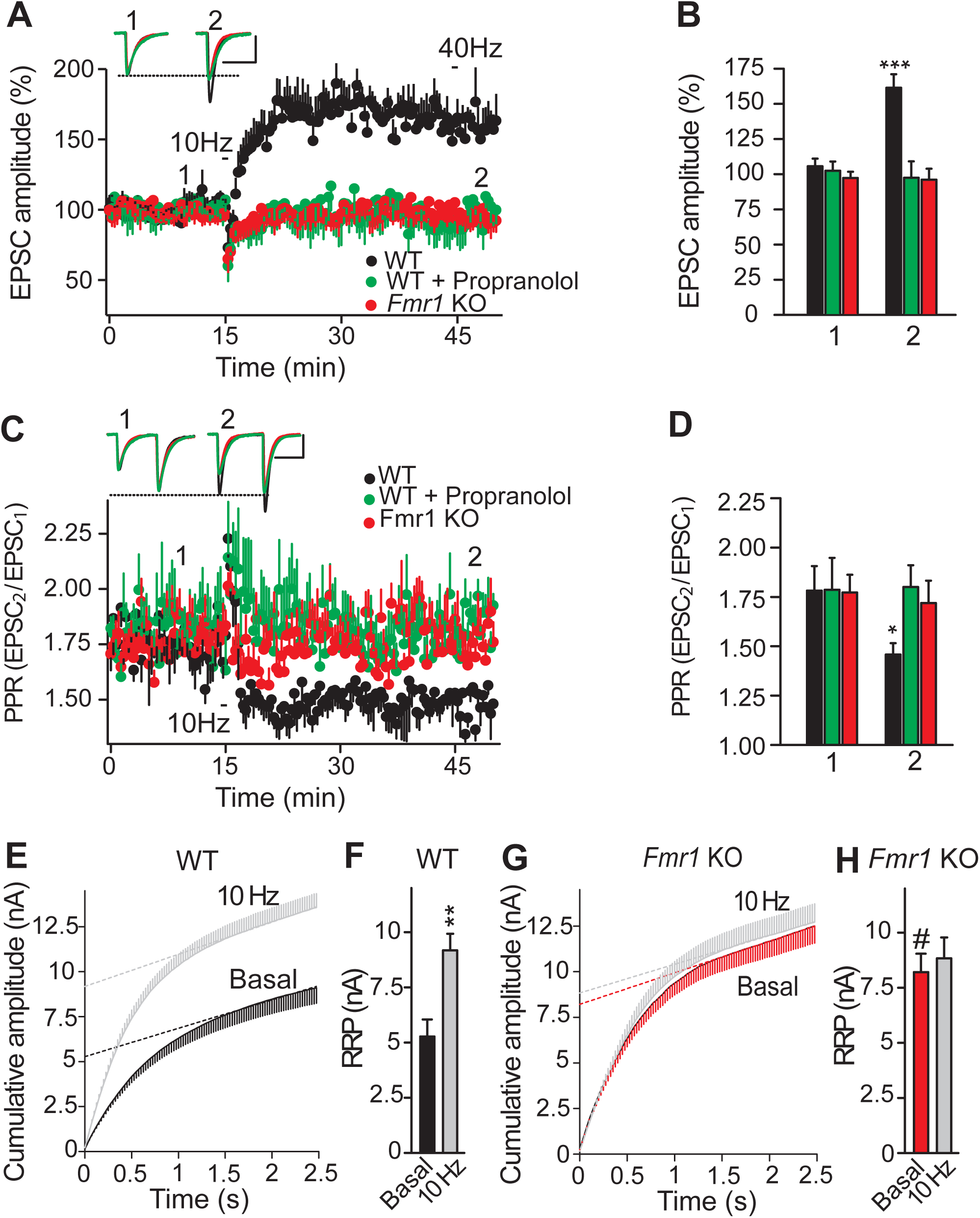
*Fmr1* KO synapses lack cerebellar PF-PC LTP. (A) Changes in EPSC amplitude induced by a 10 Hz stimulation. (B) Quantification of EPSCs 30 min after stimulation (2) compared to the respective values before stimulation (1): WT, (n=9 cells/9 slices/5 mice: *P*=0.0001, unpaired Student’s *t* test); WT/propranolol (100 μM, 30 min, n=10 cells/10 slices/6 mice: *P*=0.7125, Welch’s test); *Fmr1* KO (n=12 cells/12 slices/9 mice: *P*=0.8955, Welch’s test). (C) Changes in the PPR. (D), Quantification of the changes in PPR: WT (*P*=0.0317 Welch’s test); WT/propranolol (*P*=0.9595, unpaired Student’s *t* test); *Fmr1*KO (*P*=0.677, unpaired Student’s *t* test). (E, G), RRP size in WT and *Fmr1*KO slices. (F,H) Quantification of the RRP size: WT (n=11 cells/11 slices/5 mice: *P*=0.0015); *Fmr1*KO n=13 cells/13 slices/6 mice: *P*=0.6302). RRP size WT basal vs *Fmr1*KO basal (^#^*P*=0.0185). The data are shown as the means ± S.E.M. Scale bars in (A, H) 50 pA and 10 ms, and in (C) 25 pA and 10 ms. Unpaired Student’s *t* test in F and H. \**P*<0.05, \*\**P*<0.01, \*\*\**P*<0.001.

PF-PC LTP involves an increase in the RRP size (Martín et al., 2020) and hence, we determined if a change in the RRP size could explain the lack of LTP at *Fmr1*KO PF-PC synapses. An increase in the RRP size was evident 30 min after LTP induction at WT PF-PC synapses (*P*=0.0015, Fig. 3*E,F*). However, the basal RRP was larger in *Fmr1* KO synapses than in WT slices (*P*=0.0185) and it was not further enhanced by stimulation at 10 Hz (*P*=0.6302, Fig. 3*E,F,G,H*). As such, *Fmr1*KO synapses have a larger RRP under basal conditions that impedes LTP.

### 2.4. Reducing extracellular Ca^2+^ rescues parallel fiber LTP in Fmr1KO slices

Some presynaptic changes in *Fmr*1KO mice include a loss of functional Ca^2+^-activated K^+^ channels, which results in AP broadening during repetitive stimulation, enhanced presynaptic Ca^2+^ influx and increased neurotransmitter release (Deng et al., 2013). By activating proteins of the release machinery involved in SV docking and priming, such as Munc13 (Shin et al., 2010; Dimova et al., 2006, 2009), presynaptic Ca^2+^ can modulate the docking of SVs and the size of the RRP (Taschenberger et al., 2016; Thanawala and Regehr, 2013; Martín et al, 2018). Thus, we tested whether reducing [Ca^2+^]_e_ re-established isoproterenol mediated potentiation in *Fmr1*KO synapses. When slices were maintained at a low [Ca^2+^]_e_ concentration (1 mM), isoproterenol increased the sEPSCs amplitude (*P*=0.0021, Fig. 4*A,B*). Moreover, the aEPSC frequency of *Fmr*1KO synapses decreased to values similar to those of WT synapses in normal 2.5 mM [Ca^2+^]_e_ (*P*=0.5054, Fig. 4*D*), and consequently, exposure to isoproterenol increased the aEPSC frequency (*P*=0.0396, Fig. 4*C,D,E*) with no change in aEPSC amplitude (*P*=0.9819, Fig. 4*F,G*). When the [Ca^2+^]_e_ was lowered PF-PC LTP was also recovered at *Fmr1*KO slices (*P*=0.0001, Fig. 4*H,I)*, even though LTP was not supported in WT slices at this low [Ca^2+^]_e_ (*P*=0.9571, Fig. 4*H,I*), consistent with earlier reports on the sensitivity of PF-PC LTP to decreases in [Ca^2+^]_e_ (Myoga and Regehr, 2011). Interestingly, the rescued LTP in *Fmr1*KO slices also exhibited other features of PF-PC LTP seen in WT synapses (Martín et al., 2020), such as its sensitivity to the β-AR antagonist propranolol (*P*=0.9283, Fig. 4*H,I*) and the increase in RRP size (*P*=0.0002, Fig. 4*J,K*). Indeed, reduced [Ca^2+^]_e_ counteracted the changes that led to increased SV docking and the enhanced RRP in *Fmr1*KO PF-PC synapses, rescuing the presynaptic potentiation by β-ARs and PF-PC LTP.

**Figure 4.**
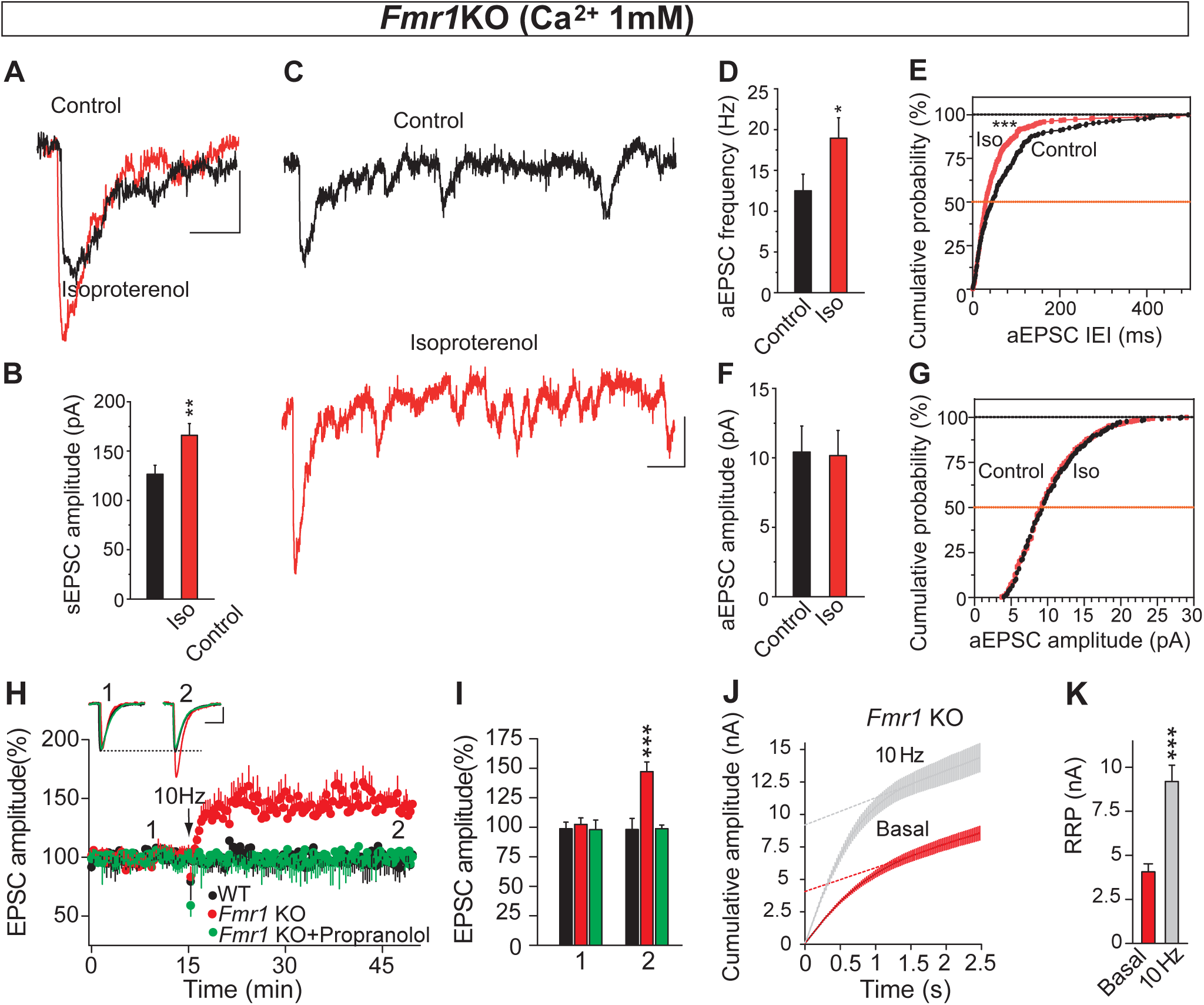
Reducing extracellular Ca^2+^ re-establishes isoproterenol induced potentiation of asynchronous release and parallel fiber LTP at *Fmr1*KO slices. Isoproterenol (100 μM, 10 min) enhances the sEPSC amplitude recorded in the presence of Sr^2+^ (1.0 mM). (B) Quantification of the effects of isoproterenol on sEPSC amplitude: (n=6: *P*=0.0021, Unpaired Student’s t test). (C) Individual traces showing asynchronous release events in control (black) and after exposure to isoproterenol (red). (D,F) Quantification of the isoproterenol induced changes in aEPSC frequency (D) (n=305 events/10 slices and n=474 events/10 slices: *P*=0.0396) and in aEPSC amplitude (F) (*P*=0.9819, unpaired Student’s t test). (E,G) Cumulative probability plots of isoproterenol induced changes in aEPSC frequency (inter event interval, IEI) (E) (*P*=0.0001) and aEPSC amplitude (G) (*P*=0.6499, Kolmogrov-Smirnov test). (H), PF LTP was recovered in *Fmr1*KO slices, yet no LTP was induced in WT slices in the presence of 1 mM Ca^2+^. Experiments in *Fmr1*KO slices were also performed in the presence of the β-AR antagonist propranolol (100 μM, added 30 min prior to LTP induction). The EPSC sample traces represent the mean of 6 consecutive EPSCs at 0.05 Hz. (I) Changes in EPSC amplitude 30 min after stimulation (2) compared to basal (1): in *Fmr1*KO (n=14 cells/14 slices/7 mice: *P*=0.0001, unpaired Student’s *t* test); in *Fmr1*KO slices treated with propranolol (n=9 cells/9 slices/4 mice: *P*=0.9283, Welch’s test); and in WT (n=10 cells/10 slices/4 mice: *P*=0.9571, unpaired Student’s *t* test). (J), RRP size in synapses from basal 10 Hz stimulated slices of *Fmr1*KO mice. (K), Quantification of the RRP in *Fmr1*KO slices (n=12 cells/12 slices/4 mice and n=12 cells/12 slices/5 mice, before and after LTP induction: \*\*\**P*=0.0002, Welch’s test). The data are the means ± S.E.M. Scale bar in (A, C) 10 pA and 100 ms; and in (H) 50 pA and 30 ms. \**P*<0.05, \*\*\**P*<0.001.

### 2.5. The mGluR4 PAM VU0155041 rescues parallel fiber LTP at Fmr1 KO slices

Since decreasing [Ca^2+^]_e_ re-establishes LTP at *Fmr1*KO synapses, it may be possible to rescue LTP using pharmacological tools that reduce Ca^2+^ influx at PF synaptic boutons, for example through the activation of G protein coupled receptors (GPCRs). Significantly, mGluR4 reduces Ca^2+^ influx and synaptic transmission at PF synaptic boutons (Pekhletski et al., 1996; Daniel and Crepel, 2001), where mGluR4 is the only group III mGluR present (Abitbol et al., 2008). When mGluR4s were activated by L-AP4 (40 μM) there was a reduction in the EPSC amplitude and once a new baseline was established, 10 Hz stimulation induced a strong and sustained increase in the EPSC amplitude (*P*<0.0001, Fig. 5*A,B*). We also tested whether VU 0155041, a potent and selective PAM of mGluR4s that is active in vivo (Niswender et al., 2008; Williams et al., 2009), rescued PF-PC LTP. VU 0155041 reduced the EPSC amplitude and after 5 min, a 10 Hz stimulation provoked a sustained increase in EPSC amplitude (*P*<0.0001, Fig. 5*A,B*). The rescued LTP at *Fmr1*KO synapses required β-AR activation, as does WT LTP, and indeed, the β-AR antagonist propranolol prevented this rescue by VU 0155041 (*P*=0.1255, Fig. 5*A,B*). VU 0155041 reduced the RRP size in *Fmr1*KO synapses under basal conditions (*P*=0.0100, Fig. 5*C,D*) and permitted a further increase by 10 Hz stimulation (*P*=0.0014, compared with VU 0155041 in basal conditions). Thus, the mGluR4 PAM VU 0155041 rescued normal RRP size and PF LTP in *Fmr1*KO slices.

**Figure 5.**
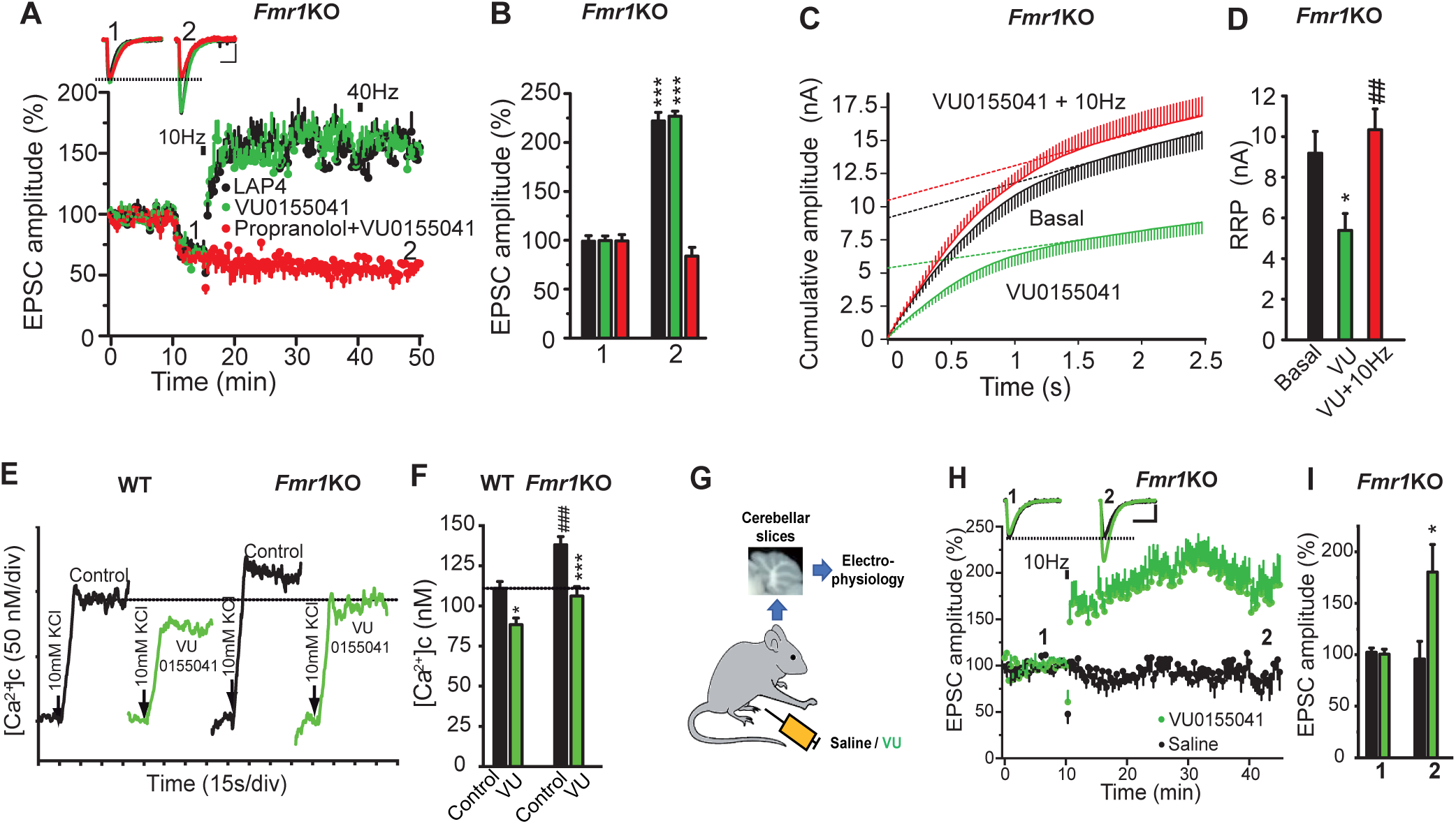
Activation of mGluR4 rescues PF-PC LTP in *Fmr1*KO slices. (A) LAP4 (40 μM) and the mGluR4 PAM VU 0155041 (100 μM) rescues PF LTP in *Fmr1*KO slices. The β-AR antagonist propranolol (100 μM) was added 30 min prior to LTP induction. EPSC traces are the mean of 6 consecutive EPSCs at 0.05 Hz. (B) Quantification of EPSC amplitude 30 min after stimulation (2) compared to the respective values before stimulation (1): LAP4 (n=10 cells/10 slices/5 mice: *P*<0.0001); VU (n=11 cells/11 slices/6 mice: *P*<0.0001); propranolol + VU (n=10 cells, 10 slices/6 mice: *P*=0.1255). (C) RRP size in *Fmr1*KO slices in the presence or absence (basal) of VU (100 μM, 10 min) and with or without 10 Hz stimulation. (D) Quantification of the RRP size in the above conditions: Basal (n=10 cells/10 slices/7 mice); VU (n=12 cells/10 slices/7 mice: *P*=0.0100); VU plus 10 Hz (n=8 cells/8 slices/4 mice: ^##^*P*=0.0014, compared to VU alone). (E) VU restores Ca^2+^ dynamics to *Fmr1*KO cerebellar synaptosomes. Changes in the cytoplasmic free Ca^2+^ concentration ([Ca^2+^]_c_) in the presence and absence of VU (100 μM) added at least 5 min prior to KCl. (F) Quantification of the changes in [Ca^2+^]_c_: KCl/WT (n=18/5 preparations); KCl/*Fmr1* KO (n=16/5 preparations: ^###^*P*<0.001 compared to KCl/WT); VU/KCl/WT (n=11/4 preparations: *P*<0.05, compared to KCl/WT); and VU/KCl/*Fmr1*KO (n=12/4 preparations: *P*<0.001, compared to KCl/*Fmr1*KO). (G) Scheme showing the preparation of cerebellar slices from VU or saline injected *Fmr1*KO mice. (H) Response to a 10 Hz stimulation in slices from VU and saline injected *Fmr1*KO mice. EPSC traces are the mean of 6 consecutive EPSCs at 0.05 Hz. (I) EPSC amplitude 30 min after stimulation (2) compared to the respective values before stimulation (1): VU (5 mg/Kg) injected *Fmr1*KO mice (n=14 cells/14 slices/10 mice, *P*=0.0117); saline injected *Fmr1*KO mice (n=10 cells/10 slices/8 mice, *P*=0.7178). The data are the means ± S.E.M. Scale bars in (A, H) 100 pA and 10 ms. Unpaired Student’s t test (B,D). Two-way ANOVA with Tukey in (F). Welch test (I). \**P*<0.05, \*\*\**P*<0.001.

Ca^2+^ homeostasis is altered in *Fmr1*KO mice (Deng et al., 2013; García-Font et al., 2019; Ferron et al., 2014), which led us to test whether VU 0155041 might rescue Ca^2+^ dynamics in *Fmr1*KO mice by measuring the depolarization induced change in the cytosolic Ca^2+^ concentrations ([Ca^2+^]_c_) of fura-2 loaded cerebellar synaptosomes. The KCl-induced increase in [Ca^2+^]_c_ was larger in *Fmr1*KO than in WT synaptosomes (*P*<0.001, Fig. 5*E,F*), compatible with the prolonged APs at *Fmr1*KO synapses (Deng et al., 2013). VU 0155041 reduced the KCl-induced change in [Ca^2+^]_c_ in WT synaptosomes (*P*<0.05, Fig.5*E,F*) and it restored the KCl-induced increase in [Ca^2+^]_c_ in *Fmr1*KO to the levels of WT synaptosomes (*P*>0.05 Fig. 5*E,F*). Together, these data indicate that Ca^2+^ homeostasis is deregulated in *Fmr1*KO cerebellar synaptosomes but can be restored with the mGluR4 PAM VU 0155041.

We also tested whether VU 0155041 injected “in vivo” rescue PF-PC LTP in cerebellar slices. *Fmr1*KO mice were injected (i.p.) with VU 0155041 (or the saline vehicle alone) and cerebellar slices were prepared 2 hours later. PF-PC LTP was rescued in slices from VU 0155041 injected *Fmr1*KO mice (*P*=0.0117, Fig. 5*G,H,I*) but not in those from *Fmr1*KO mice injected with saline alone (*P*=0.7178, Fig. 5*G,H,I*). Then, the injection of mGluR4 PAM VU 0155041 “in vivo” also rescued PF-PC LTP in cerebellar slices.

### 2.6. VU0155041 ameliorates the motor learning and social deficits of Fmr1KO mice

In order to assess the behavioral consequences of the loss of PF-PC LTP we tested the performance of *Fmr1*KO mice in tests that evaluate motor learning. *Fmr1* KO mice display impaired motor learning in a forelimb-reaching task (Padmashri et al., 2013). In this test, mice are trained to use their forelimbs to grasp and retrieve food pellets through a narrow slit (Fig. 6*A*), and the cerebellum contributes substantially to the coordination of the skilled movements that require speed, smoothness and precision, such as reaching to grasp movements (Sakayori et al., 2019; Becker and Person, 2019). We tested whether rescuing the PF-PC LTP with VU 0155041 improved skilled reaching. After two days of habituation, the animal’s efficiency (number of pellets retrieved/number of attempts) was measured over 5 days and compared to that on day 1. After 5 days, WT mice that received saline (WT sal) increased test performance (*P*<0.0001), unlike similarly treated *Fmr1*KO mice (*Fmr1* KO sal) (*P*=0.244, Fig. 6*B*). Interestingly, VU 0155041 injection improved the learning of this task by *Fmr1*KO mice (*Fmr1*KO VU) (*P*=0.004, Fig. 6*B*), while it did not alter that of WT mice (WT VU) (*P*<0.0001, Fig. 6*B*).

**Figure 6.**
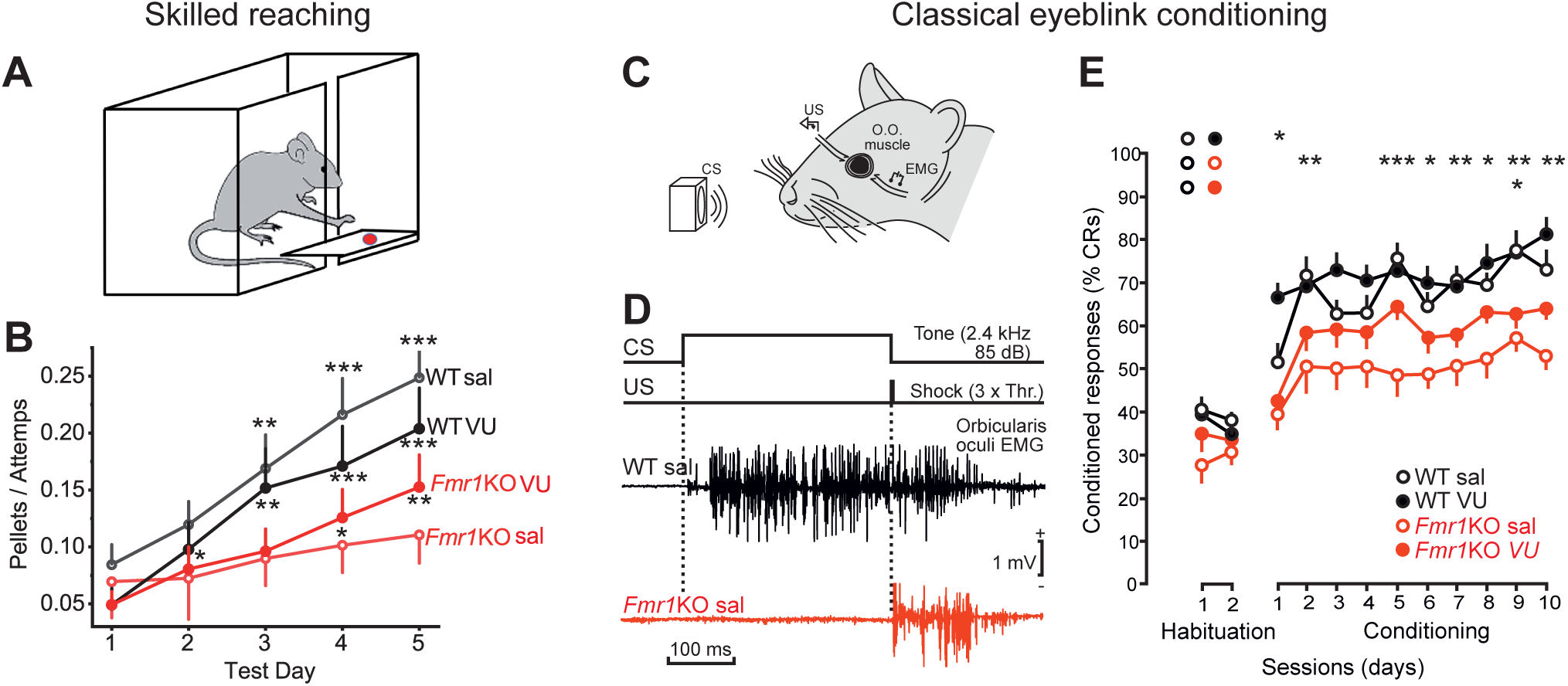
VU 0155041 ameliorates skilled reaching and classical eyeblink conditioning deficits of *Fmr1*KO mice. (A) Skilled reaching test. Mice use their forelimbs to grasp and retrieve food pellets through a narrow slit. (B) Efficiency in test performance (number of pellets retrieved per attempt) in the four experimental groups: WT/sal (n=31); *Fmr1*KO sal (n=33); WT VU (n=31); and *Fmr1*KO VU (n=32). The increase in efficiency is shown for day 2, 3, 4 and 5 relatives to day 1: WT sal: *P*=0.078, *P*=0.009, *P*<0.0001 and *P*<0.0001; *Fmr1* KO sal: *P*=0.883, *P*=0.515, *P*=0.319 and *P*=0.244; WT VU: *P*=0.015, *P*=0.002. *P*<0.0001 and *P*<0.0001; *Fmr1* KO/VU: *P*=0.112, *P*=0.137, *P*=0.02 and *P*=0.004. Two-way repeated measures ANOVA followed by LSD (C) Classical eyeblink conditioning was evoked with a conditioning stimulus (CS) consisting of a 350 ms tone (2.4 kHz, 85 dB) supplied by a loudspeaker located 50 cm in front of the animal’s head. The unconditioned stimulus (US) was presented at the end of the CS, and consisted of an electrical shock (a square, cathodal pulse, lasting for 0.5 ms) presented to the left supraorbital nerve. Conditioned responses (CRs) were determined from the EMG activity of the orbicularis oculi (O.O.) muscle ipsilateral to US presentations. (D) Examples of EMG recordings collected from representative WT sal and *Fmr*1KO sal mice during the 8th conditioning session. Note the presence of a noticeable CR in the WT sal mouse and its absence in the Fmr1KO sal animal. (E) CRs after 10 conditioning sessions of WT sal (n=10), *Fmr1*KO sal (n=10), *P*<0.01, WT VU (n=10), *P*>0.05 and *Fmr1*KO VU (n=10), *P*>0.05. The data are the mean ± S.E.M. (\**P*<0,05; \*\**P*<0.01; \*\*\**P*<0.001). Two-way repeated measures ANOVA followed by Holm-Sidak method (all comparisons were to WT sal).

We also tested classical eyeblink conditioning and the VOR, two paradigms that specifically evaluate cerebellar-dependent motor learning related to plasticity at PF-PC synapses (Ito, 2002; Gutierrez-Castellanos et al., 2017; Hirano, 2018). *Fmr1*KO mice show deficits in classical eyeblink conditioning (Koekkoek et al., 2005). In the classical eyeblink conditioning the mouse learn to associate a conditioned stimulus (CS), such as a tone, with an unconditioned stimulus (US) such as a mild electric shock to the supraorbital nerve, which evokes eyeblinks (Fig. 6*C,D*). As a result of the CS-UC association during training the eyeblink conditioned response (CR) is progressively enhanced (Sánchez-Campusano et al., 2009). After 10 conditioning sessions there was a significant deficit in the number of CRs of *Fmr1*KO sal mice compared to WT sal mice (*P*<0.01, Fig. 6*E*). VU 0155041 administration improves the performance of the *Fmr1*KO (*P*=0.072, Fig. 6*E*), but not that of the WT (*P*=0.333, Fig. 6*E*).

The VOR helps to stabilize gaze when the head turns (Fig. 7A). The cerebellum plays an important role in the control of phase and gain dynamics of the VOR (Ito, 2002). Illustrative examples of eye movements during table rotation at different frequencies are shown (Fig. 7*A*). *Fmr1*KO sal mice showed significant deficits in gain (0.1 Hz: *P*<0.05, Fig. 7*B*) and phase (0.6 Hz: *P*=0.016, Fig. 7*C*). VU 0155041 administration compensated the differences in gain (0.1 Hz: *P*>0.05) (Fig 7*B*), and phase (*P*>0.446, Fig. 7*C*). *Fmr1*KO sal mice presented an increase in the number of fast phases per vestibuloocular cycle (0.6 Hz: *P*<0.05; 0.3 Hz: *P*=0.013), that was compensated following VU 0155041 administration (0.6 Hz: *P*>0.05; 0.3 Hz: *P*=0.470, Fig. 7*D*). The relation between vestibuloocular gain and fast phases frequency was also different between WT sal and *Fmr1*KO sal mice (0.6 Hz: *P*<0.05; 0.3 Hz: *P*<0.01, Fig. 7*E*), and these differences were restored after VU 0155041 administration to *Fmr1*KO (0.6 Hz: *P*>0.05; 0.3 Hz: *P*>0.05, Fig. 7*E*). Then, VU 0155041 may offer some therapeutic relief to the motor learning deficits in FXS.

**Figure 7.**
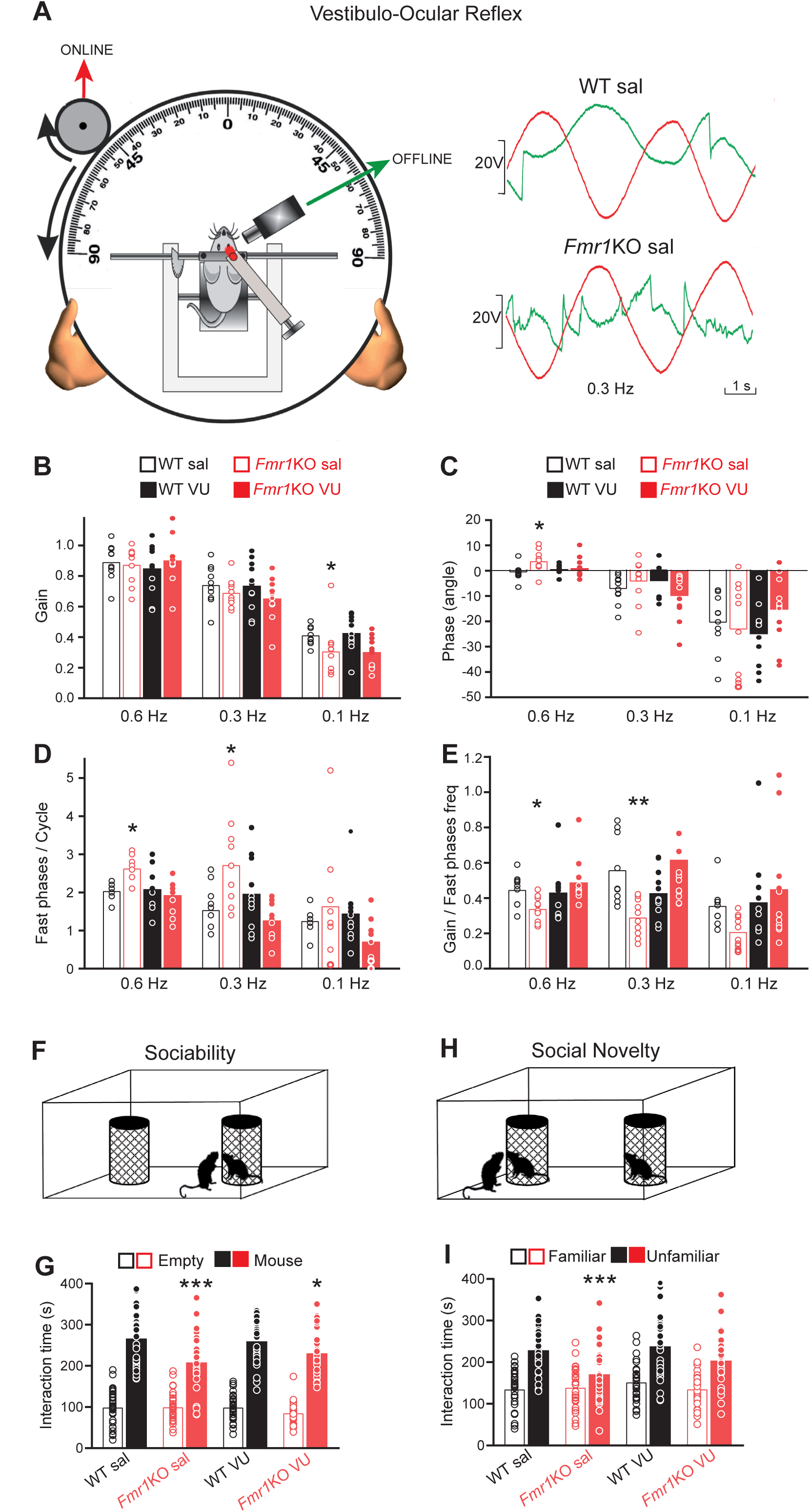
VU 0155041 ameliorates VOR and social interaction deficits of *Fmr1*KO mice. A, Experimental design. Table rotations (red) and eye positions (green). Examples of eye movements. (B) VOR gain. 0.1 Hz: WT sal (n=10), *Fmr1*KO sal (n=10), *P*<0.05, WT VU (n=11), *P*>0.05 and *Fmr1*KO VU (n=11), *P*>0,05. Kruskal-Wallis followed by Dunn’s test. (C) VOR phase. 0.6 Hz: *Fmr1*KO sal, *P*=0.016, WT VU P=0.382, *Fmr1*KO VU, *P*=0.446. One-way ANOVA followed by Holm-Sidak. (D) Fast phases per vestibuloocular cycle. 0.6 Hz: *Fmr1*KO sal, *P*<0.05, WT VU *P*>0.05 and *Fmr1*KO VU, *P*>0.05, Kruskal-Wallis followed by Dunn’s test. 0.3 Hz: *Fmr1*KO sal, *P*=0.013, WT VU, *P*=0.468, *Fmr1*KO VU, *P*=0.470, one-way ANOVA followed by Holm-Sidak. (E), Relation between VOR gain and fast phases frequency. 0.6 Hz: *Fmr1*KO sal, P<0.05, WT VU, *P*>0.05, *Fmr1*KO VU, *P*>0.05. Kruskal-Wallis followed by Dunn’s method. 0.3 Hz: *Fmr1*KO sal, *P*<0.01, WT VU, *P*>0.05, *Fmr1*KO VU, *P*>0.05, Kruskal-Wallis followed by Dunn’s method. (F) Sociability test set-up. (G) the interaction time (s) with the mouse-containing and empty cages is measured: WT sal (n=31); *Fmr1*KO sal (n=30, *P*<0.0001); WT VU (n=31, *P*=0.652); (*Fmr1*KO VU (n=32, *P*=0.021). (H), Social novelty test set-up. (I), The social novelty test measures the interaction time with familiar and unfamiliar mice: WT sal (n=31); *Fmr1* KO sal (n=30, *P*<0.001); WT VU (n=31, *P*=0.578); (*Fmr1* KO VU (n=32, *P*=0.131). Two-way repeated measures ANOVA followed by LSD (G,I). The data are the mean ± S.E.M. (\**P*<0.05; \*\**P*<0.01, \*\*\**P*<0.001). All post-hoc comparisons were to WT sal.

*Fmr1*KO mice shown no defects in other motor tests such as the rotarod, that measures the time that each animal remained on the rod of and an accelerating rotarod treadmill (time to fall) (*Supplementary Material:* Fig. S5*A*), or in the elevated path that measures the time spent by a mouse placed in the center on an elevated bar to reach one of the two platforms (*Supplementary Material:* Fig. S5*B*)

The cerebellum also controls some non-motor activities such as social cognition (Van Overwalle et al., 2020). Thus, we assessed whether VU 0155041 also ameliorates the social impairments of these *Fmr1*KO mice. The sociability test evaluates the time animals interact with a cage containing another mouse as opposed to that with an empty cage (Fig 7*F*). *Fmr1*KO sal mice spend less time than WT sal mice interacting with another mouse in a cage (*P*<0.0001), although VU 0155041 injection partially reduced this deficit (*P*=0.0210, Fig. 7*G*). We also evaluated social novelty by measuring the interaction time with a familiar as opposed to an unfamiliar mouse (Fig. 7*H*). *Fmr1*KO sal mice spend less time with unfamiliar mice than WT sal mice (*P*=0.001), yet again this deficit was reversed by the injection of VU 0155041 (*P*=0.131, Fig. 7*I*). These results indicate that VU 0155041 ameliorates the alterations in social behavior of *Fmr1*KO mice.

## 3. DISCUSSION

In this study, the lack of presynaptic PF-PC LTP in *Fmr1*KO mice is shown to be a consequence of the increase in SV docking and priming, and in the RRP size, which precludes further potentiation of neurotransmitter release. These parameters can be restored by either diminishing the [Ca^2+^]_e_ concentration or through a decrease in Ca^2+^ influx at PFs induced by the mGluR4 PAM, VU 0155041. *Fmr1*KO mice also show deficits in skilled reaching, classical eyeblink conditioning and in the VOR. Interestingly, administration of VU 0155041 ameliorates these motor learning deficits and the impaired social interactions of these *Fmr1*KO mice.

The RRP of SVs is made up of docked and fully primed SVs, whose membrane fusion can be triggered by Ca^2+^ influx. The number of docked SVs is strongly related to the size of the RRP (Rosenmund and Stevens, 1996; Schikorski and Stevens, 2001) and therefore, an increase in tightly docked and fully primed SVs may explain the increased aEPSC frequency at *Fmr1*KO synapses. However, it is important to understand why *Fmr1* KO synapses have more docked/primed SVs in order to design strategies to revert this phenotype. Two protein interactions influence the docking and priming of SVs: the formation of the Munc13-Rim1-Rab3 complex (Dulubova et al., 2005); and the assembly of the SNARE complex promoted by the priming activity of Munc13 (Ma et al., 2011). As such, neurons deficient in Munc13s have no docked SVs (Imig et al., 2014) and Munc13-2 deficient synapses fail to display the mGluR7 receptor mediated potentiation associated to enhanced SV docking/priming (Martín et al., 2018). Munc13-1 is translocated to membranes upon DAG binding (Brose et al, 1995; Brose and Rosenmund, 2002), promoting its interaction with RIM1 to form a heterodimer with priming activity (Deng et al., 2011b). In addition to the DAG binding C1 domain (Betz et al., 2001), Munc13-1 is also activated by Ca^2+^ through its Ca^2+^-CaM (Dimova et al., 2006, 2009) and Ca^2+^-phospholipid binding domains (Shin et al., 2010). Cerebellar granule cells express Munc13-1 and Munc13-3, the latter also containing binding sites for Ca^2+^ and DAG (Augustin et al., 2001). Thus, it is likely that the decrease in [Ca^2+^]_e_ dampens Munc13 activity and that this provokes a reduction in the RRP size of sufficient magnitude to counterbalance the effect that elevated Ca^2+^ and DAG levels may have in *Fmr1* KO synapses (Deng et al., 2013; Tabet et al., 2016).

Decreasing [Ca^2+^]_e_ re-establishes β-AR mediated potentiation and PF-PC LTP at *Fmr1*KO synapses. However, as this strategy has limited translational potential, we tested whether PF-PC LTP can be also rescued pharmacologically by activating presynaptic responses that reduce Ca^2+^ influx at PF synaptic boutons. We found that mGluR4 activation, either by the group III mGluR agonist L-AP4 or the mGluR4 specific PAM VU 0155041, restores PF-PC LTP. This response is consistent with the presynaptic localization of these receptors at PF-PC synapses (Kinoshita et al., 1996), and with their capacity to depress synaptic transmission and presynaptic Ca^2+^ influx (Daniel and Crepel, 2001). VU 0155041 also restores the Ca^2+^ responses of cerebellar synaptosomes to those of WT synaptosomes.

We found that the acquisition of motor skills that affect movements involved in reaching and grasping is impaired in *Fmr1*KO mice as shown previously by (Padmashri et al., 2013), consistent with the deficient fine motor skills acquisition of FXS patients (Will et al., 2018). The motor skills required for reaching and grasping are high-level functions that involve multiple circuits and brain regions, including the cerebellum, basal ganglia and motor cortex (Shmuelof and Krakauer, 2011). The cerebellum contributes substantially to the coordination of skilled movements, such as those involved in reaching to grasp tasks (Sakayori et al., 2019; Becker and Person, 2019). Thus, patients with lesions in the cerebellar cortex have impaired reach to grasp movements (Rand et al., 2000). At the cellular level, PCs encode limb movements during reaching tasks (Sakayori et al., 2019). It is well established that classical eyeblink conditioning and VOR represent two forms of cerebellar motor learning. For many years the prevailing view has been that cerebellar motor learning depends on postsynaptic LTD at the parallel fiber to Purkinje cell synapses (Ito, 2001). However, it has also been shown that postsynaptic LTP also contributes to cerebellar motor learning providing a reversal of the LTD induced synaptic changes (Gutierrez-Castellanos et al., 2017). Now we found that cerebellar motor learning may also depend on presynaptic PF-PC LTP, thus widening the dynamic range of the synaptic changes that control this cerebellar function.

In addition to the lack of presynaptic PF-PC LTP described here, other changes at PF-PC *Fmr1* KO synapses include spine elongation and enhanced postsynaptic LTD (Koekkoek et al., 2005), both of which are associated with motor learning deficits (Galliano et al., 2013). It is therefore attractive to hypothesize that the presynaptic changes at *Fmr1*KO PF synapses described here (enhanced SV docking, RRP size and aEPSC frequency) could represent a compensatory mechanism to counterbalance the postsynaptic changes (enlarged and immature spines, and enhanced LTD) (Koekkoek et al., 2005). Interestingly, presynaptic LTP in the anterior cingulate cortex is also abolished in *Fmr1*KO mice (Koga et al., 2015).

We also found that rescuing PF LTP with the mGluR4 PAM, VU 015541 favors the social interaction of *Fmr1*KO mice. Recent work revealed that the cerebellum controls not only motor functions but also social behavior (Van Overwalle et al., 2020). Early developmental damage to the cerebellum is associated with deficits in social contact in autism (Limperopoulos et al., 2007), and cerebellar abnormalities that affect the structure and function of PCs have been detected in mouse models of autism (Tsai et al., 2012), linking cerebellar dysfunction with autistic behavior. FXS patients also suffer cerebellar alterations, with a reduced size and density of PCs (Greco et al., 2011).

In conclusion, our data points to the cerebellum as a potential target for pharmacological intervention to ameliorate not only the motor learning deficits but also, the social interactions of FXS patients. We are aware that one limitation of the study is that activating mGluR4s via systemic administration of VU 155041 can have broad effects that are likely to affect many neurons and circuits across several brain regions. However, cerebellar granule cells have the strongest expression of mGluR4 in the brain (Corti et al., 2002) and therefore, it is likely to be in this region where the effects of VU 155041 would be most significant.

## 4. MATERIALS AND METHODS

### 4.1. Synaptosome preparation

*Fmr1*KO mice or WT littermate (3 months old) were anaesthetized with isoflurane (1.5-2% in a mixture of 80% synthetic air/20% oxygen) and sacrificed by decapitation. Synaptosomes were purified from 4-5 cerebella on discontinuous Percoll gradients (GE Healthcare, Uppsala, Sweden). Briefly, the tissue was homogenized and centrifuged for 2 min at 2,000 x g, and the supernatant was then centrifuged again for 12 min at 9,500 x g. The pellets obtained were gently resuspended in 0.32 M sucrose [pH 7.4] and placed onto a 3 ml Percoll discontinuous gradient containing: 0.32 M sucrose; 1 mM EDTA; 0.25 mM DL-dithiothreitol; and 3, 10 or 23 % Percoll [pH 7.4]. After centrifugation at 25,000 x g for 10 min at 4 ºC, the synaptosomes were recovered from between the 10 % and 23 % Percoll bands, and they were resuspended in HEPES buffered medium (HBM) (Millán et al., 2002). The preparation of cerebellar synaptosomes largely represents the synaptic boutons of granular cells, by far the most abundant cells in the brain (Andersen et al., 1992).

### 4.1.1. Glutamate release

Glutamate release from cerebellar synaptosomes was assayed by on-line fluorimetry (1) based on the reduction of NADP^+^ (1 mM: Calbiochem) by glutamate dehydrogenase (Sigma-Aldrich, St. Louis, MO, USA). The fluorescence of the NADPH generated was measured in a Perkin Elmer LS-50 luminescence spectrometer at excitation and emission wavelengths of 340 and 460 nm, respectively, and using FL WinLab v. 4.00.02 software. Spontaneous glutamate release was determined in the presence of the Na^+^-channel blocker, Tetrodotoxin (TTx, 1 μM: Abcam, Cambridge, UK).

### 4.1.2. cAMP accumulation

The accumulation of cAMP was determined using a cAMP dynamic 2 kit (Cisbio, Bioassays, Bagnols sur-Cèze, France). Synaptosomes were incubated for 1 h at 37 ºC (0.67 mg/ml) in HBM and after 15 min, 1.33 mM Ca^2+^ and 1 mM of the cAMP phosphodiesterase inhibitor 3-isobutyl-1-methylxanthine (IBMX: Calbiochem, Damstard, Germany) were added to the synaptosomes for 15 min. Isoproterenol (100 μM) was then added for 10 min, the synaptosomes were collected by centrifugation and transferred to a 96-well assay plate, and the assay components were added having been diluted in lysis buffer: the europium cryptate-labeled anti cAMP antibody and the d2-labeled cAMP analog. After incubation for 1 h at room temperature (RT), the europium cryptate fluorescence and TR-FRET signals were measured over 50 ms on a FluoStar Omega microplate reader (BMG Lab Technologies, Offenburg, Germany) at 620 and 665 nm, respectively, after excitation at 337 nm [see (García-Font et al., 2019)]. The data were obtained using Omega BMG Labtech v.1.00 software.

### 4.1.3. Immunofluorescence

Immunofluorescence was performed using an affinity-purified rabbit polyclonal antiserum against β_1_-AR (1:200, Cat# sc-568: Santa Cruz Biotechnology, RRID:AB_2225388) and a mouse monoclonal antibody against synaptophysin 1 (1:500, Cat# 101 011: Synaptic Systems, RRID:AB_887822), as described previously (Ferrero et al., 2016). As a control for the immunofluorescence, the primary antibodies were omitted from the staining procedure, whereupon no immunoreactivity resembling that obtained with the specific antibodies was evident. After washing in Tris Buffered Saline, TBS, the labelled synaptosomes were incubated for 2 hours with secondary antibodies diluted in Tris TBS: Alexa fluor 488 Donkey anti-mouse IgG (1:500, Cat# A-21202: Invitrogen, RRID:AB_141607) and Alexa fluor 594 Donkey anti-rabbit IgG (1:500, Cat# A-21207: Invitrogen, RRID:AB_141637). After several washes in TBS, the coverslips were mounted in Prolong Antifade Kit (Molecular Probes, Eugene, OR, USA), and the synaptosomes were viewed on a Nikon Diaphot microscope equipped with a 100x objective, a mercury lamp light source and fluorescein-rhodamine Nikon filter sets. Images were acquired with AQM 4800_80 software and analyzed using Image J 1.43m software.

### 4.1.4. Western Blotting

The P2 crude synaptosomal fraction (4 μg of protein per lane) was diluted in Laemmli loading buffer with β-mercaptoethanol (5% v/v), resolved by SDS-PAGE (8% acrylamide, Bio-Rad), and analyzed in Western blots following standard procedures. The proteins were transferred to PVDF membranes (Hybond ECL: GE Healthcare Life Sciences, Madrid, Spain) and after several washes, the membranes were probed with a polyclonal rabbit anti-β1-AR antiserum diluted 1:200 (Santa Cruz Biotechnology Cat# sc-568, RRID:AB_2225388) and a monoclonal mouse anti-β-actin antibody diluted 1:5000 (Sigma cat# A2228, RRID:AB_476697). After several washes, the membranes were incubated with the corresponding IRD-labeled secondary antibodies: goat anti-rabbit and goat anti-mouse coupled to IRDye 800 (LI-COR Biosciences Cat# 925-32211, RRID:AB_2651127) or IRDye 680 (LI-COR Biosciences Cat# 925-68020, RRID:AB_2687826). The membranes were scanned in an Odyssey Infrared imaging system, and the immunolabeling of proteins was compared by densitometry and quantified using Odyssey 2.0 software. The data were normalized to the β-actin signal to account for loading differences.

### 4.1.5. Cytosolic free Ca^2+^

The cytosolic free Ca^2+^ concentration ([Ca^2+^]_c_) was measured with fura-2. P2 crude synaptosomes were resuspended in HBM (1.5 mg/ml) with 16 μM BSA in the presence of CaCl_2_ (1.3mM) and fura-2-acetoxymethyl ester (fura 2-AM, 5 μM: Molecular Probes, Eugene, OR, USA), and they were incubated at 37 °C for 25 min. After fura-2 loading, the synaptosomes were pelleted and resuspended in 1.1 ml fresh HBM medium without BSA. A 1 ml aliquot was transferred to a stirred cuvette and CaCl_2_ (1.3 mM) was added. Fluorescence was monitored at 340 and 510 nm, taking data points at 0.3 s intervals, and the [Ca^2+^]_c_ was calculated using the equations described previously (Grynkiewicz et al., 1985).

### 4.2. Electrophysiology

*Fmr1* KO mice or WT littermate (18-30 days old) were anaesthetized with isoflurane (1.5-2% in a mixture of 80% synthetic air/20% oxygen) and sacrifice by decapitation. Cerebellar parasagittal slices (325 μm thick) were obtained in ice-cold Ringer’s solution [119 mM NaCl, 2.5 mM KCl, 1.3 mM MgSO_4_, 2.5 mM CaCl_2_, 26 mM NaHCO_3_, 1 mM NaH_2_PO_4_, 10 mM glucose] on a Leica VT 1200S vibratome. The slices were kept in a holding chamber containing Ringer’s solution for at least 1 hour and then transferred to a superfusion chamber for recording. The Ringer’s solution in the superfusion chamber was supplemented with 0.1 mM picrotoxin to block the GABA_A_ receptors and it was bubbled with 95% O_2_/5% CO_2_ at a flow rate of 1 mL/min. Recordings of the PF-PC synapses were obtained at 25 ºC using a temperature controller (TC-324C Warner-Instruments) as described previously (Salin et al., 1996).

Theta capillaries with a 2–5 μm tip and filled with Ringer’s solution were used for bipolar stimulation. The electrodes were connected to a stimulator (S38, GRASS) through an isolation unit and placed near the pial surface of the molecular layer to stimulate PF input to PCs. Stimuli (<50 pA, 100 ms) were delivered at 0.05 Hz with paired pulses applied 80 ms apart to obtain the PPR as EPSC_2_/EPSC_1_.

Whole cell recordings from individual PCs were obtained with a PC-ONE amplifier under voltage-clamp conditions and the membrane potential was held at -70 mV to record glutamatergic evoked EPSCs (eEPSCs). Signals were fed to a Pentium-based PC through a DigiData1322A interface board (Axon Instruments) and the pCLAMP 10.2 software was used to generate stimuli, as well as for data display, acquisition, storage and analysis. Patch pipettes (3-4 MΩ) were pulled from thin-walled borosilicate glass (1.5 mm outer diameter) on a P-97 puller (Sutter-Instrument), and they were filled with an internal solution containing: 122.5 mM cesium gluconate, 10 mM HEPES, 10 mM BAPTA, 2 mM Mg-ATP, 8 mM NaCl and 5 mM QX-314-Br (pH 7.3 adjusted with CsOH, osmolarity 290 mOsm). Series and input resistances were monitored throughout the experiment using a -5 mV pulse, and the recordings were considered stable when the series and input resistances, resting membrane potential and stimulus artifact duration did not change >20%. Cells that did not meet these criteria were discarded. To avoid irreversible effects of agonists/antagonists/inhibitors or LTP protocols, only one neuron per slice was analyzed. Recordings were also made in cerebellar slices from *Fmr1* KO mice 2 hours after intraperitoneal injection of the mGluR4 PAM, VU 0155041 (5 mg/Kg) (Duvoisin et al., 2011) or the saline vehicle alone.

To measure aEPSCs, CaCl_2_ was replaced with 2.5 mM SrCl_2_, the stimuli were adjusted to yield a EPSC amplitude between 150 and 250 pA, and they were delivered every 20 s. Asynchronous release associated with each stimulus was estimated after 20 ms and over 500 ms. The asynchronous events associated with the last six stimuli of a 5 min period were quantified in the basal condition and after 10 min in the presence of isoproterenol (100 μM).

For LTP induction, a tetanic train of 100 stimuli delivered at 10 Hz was applied at least 15 minutes after the beginning of the recording to permit adequate time for the diffusion of BAPTA into the dendritic tree. The baseline PF eEPSC amplitude was less than 300 pA to avoid sodium spikes that escaped voltage clamp, particularly during and after tetanization.

The size of the RRP was estimated as described previously (Schneggenburger et al., 2002). To minimize the variability of these estimates, the stimulus intensity prior to LTP induction was adjusted to yield EPSC amplitudes between 150-200 pA. A tetanic train (100 stimuli at 40 Hz) was applied 30 minutes after application of the 10 Hz train to induce presynaptic LTP. The cumulative EPSC amplitudes during this train were plotted and the y-intercept that extended from the linear part of the curve (times longer than 1.5 s, when the cumulative amplitude curve reaches a steady state) was used to estimate the size of the RRP. When the effect of decreasing [Ca^2+^]e on RRP size was studied, the slices were maintained at low Ca^2+^ for at least 1 h and they were incubated with 2.5 mM Ca^2+^ for 3 min prior to applying the tetanic train. OriginLab 8 software was used for plotting and fitting.

### 4.3. Electron microscopy analysis of synaptic vesicle distribution at the active zone

Parasagittal slices (325 μm thick) from the cerebellum were obtained as described above for the electrophysiology experiments, transferred to an immersion recording chamber and superfused at 1 mL min^−1^ with gassed Ringer’s solution including 0.1 mM picrotoxin. In some cases, the slices were also treated with isoproterenol (100 μM) for 10 min. The slices were fixed immediately afterwards by immersion in 3.5% glutaraldehyde in 0.1 M PB (pH 7.4) at 37 °C for 45 min and they were then left in glutaraldehyde solution for 30 min at RT before storing them for 20 h at 4 °C. The slices were then rinsed six times with large volumes of 0.1 M Phosphate Buffer (PB) and post-fixing it in 1% OsO4–1.5% K_3_Fe(CN)_6_ for 1 h at RT. After dehydrating through a graded series of ethanol (30, 50, 70, 80, 90, 95 and 100%), the samples were then embedded using the SPURR embedding kit (TAAB, Aldermaston, UK). Ultrathin ultramicrotome sections (70–80 nm thick: Leica EM UC6 Leica Microsystems, Wetzlar, Germany) were routinely stained with uranyl acetate and lead citrate, and images were obtained on a Jeol 1010 transmission electron microscope (Jeol, Tokyo, Japan). Randomly chosen areas of the cerebellar molecular layer were then photographed at 80,000x magnification and only asymmetric synapses with clearly identifiable electron-dense postsynaptic densities were analyzed with ImageJ software. The number of SVs was determined by measuring the distance between the outer layer of the vesicle and the inner layer of the AZ membrane, and distributed in 10 nm bins. The SVs that were at the maximal distance of 10 nm were considered as docked vesicles. SVs were also distributed in 5 nm bins to distinguish fully primed and tightly docked SVs (0-5 nm) from loosely docked SVs (5-10 nm). The data were analyzed blind to the genotype and treatment, and the images were analyzed using Image J 1.43m and Origin 8.0 software.

### 4.4. Drug application in “in vivo” experiments

“In vivo” experiments were carried out with 3-month-old male *Fmr1*KO mice and littermate WT. Animal were housed in individual cages until the end of the experiment. Mice were kept on a 12-h light/dark cycle with constant ambient temperature (22 ± 0.5°C) and humidity (55 ± 3%). They had food and water available ad libitum.

The mice were intraperitoneally (i.p.) injected with mGluR4 PAM (VU 0155041, 5 mg/Kg) (Duvoisin et al., 2011), or the saline vehicle alone, 2 hours prior to performing the behavioral experiments. Four experimental groups were established: WT and *Fmr1*KO mice injected with either saline or VU 0155041. Animals from each genotype were randomly allocated to one of two groups (saline or VU 0155041).

#### 4.4.1. Rotarod

We used an accelerating rotarod treadmill (Ugo Basile, Varese, Italy). A mouse was placed on the rod and tested at 2–20 rpm (of increasing speed) the first day, for a maximum of 5 min. Mice were tested two times with an interval of one hour during the same first session. Animals were re-tested 24 and 48 h later. Between these trials, mice were allowed to recover in their cages. The total time that each animal remained on the rod was computed as latency to fall (s), recorded automatically by a trip switch located under the floor of each rotating drum. Results were evaluated by averaging the data collected from each of the 4 trials (Rossi et al., 2020).

#### 4.4.2. Elevated path

The elevated path consisted of a 40 cm long, 5 cm wide bar located 60 cm over a soft cushion. Each mouse was placed in the center of the elevated bar and allowed a maximum of 40 s to reach one of the platforms (12 × 12 cm) located at each end of the bar. The time spent to reach one of the two platforms was quantified (Rossi et al., 2020).

#### 4.4.3. Skilled reaching test

Mice were food deprived (70% of normal intake) before performing the test (Tomassy et al., 2010). Briefly, a Plexiglas reaching box was used (20 cm long × 8 cm wide × 20 cm high) with a 1 cm wide vertical slit in the front of the box. Animals have to reach the palatable food pellet (20 mg dustless precision sucrose-flavored food pellets, F0071: Bio Serv) on a shelf (4 cm wide × 8 cm long) in front of the vertical slit. There is a 4 mm gap between the platform that holds the food pellets and the slit that prevents the mice from sliding the food pellets toward them. The mice were habituated to the food pellets by during the two days before testing (20 min each). Those animals that did not even attempt to grasp the pellet were discarded from the study. In the first day the food pellets were put in the box, and during the second day on the shelf. The 5 days of testing consisted of a 20 min session each day. Pellet grasping and retrieval was scored as a success, whereas pellet displacement without retrieval or grasping the pellet, dropping it before it was retrieved, and pellet displacement were considered an unsuccessful attempt. Results were represented as the ratio of total successes/total attempts.

#### 4.4.4. Animal’s preparation for chronic recording experiments

Animals were deeply anesthetized with 1.2% isoflurane supplied from a calibrated Fluotec 5 (Fluotec-Ohmeda, Tewksbury, MA, USA) vaporizer at a flow rate of 0.8 L/min oxygen. The gas was delivered via an anesthesia mask adapted for mice (David Kopf Instruments, Tujunga, CA, USA). Animals were implanted with bipolar recording electrodes in the left orbicularis oculi muscle and with bipolar stimulating electrodes on the ipsilateral supraorbital nerve. Implanted electrodes were made of 50 μm, Teflon-coated, annealed stainless steel wire (A-M Systems, Carlsborg, WA, USA), with their tips bent as a hook to facilitate a stable insertion in the orbicularis oculi muscle and bared of the isolating cover for 0.5 mm. Two 0.1-mm bare silver wires were affixed to the skull as ground. The 6 wires were soldered to a six-pin socket and the socket fixed to the skull with the help of 2 small screws and dental cement. In a final surgical step, a holding system was fixed to the skull for its proper stabilization during head rotation and eye movement recordings. Further details of this chronic preparation have been detailed elsewhere (Gruart et al., 2006; Sergaki et al., 2017).

#### 4.4.5. Classical eyeblink conditioning

Experimental sessions started a week after surgery and were carried out with six animals at a time. Animals were placed in individual and ventilated plastic chambers (5 × 5 × 10 cm) located inside a larger Faraday box (35 × 35 × 25 cm). For classical eyeblink conditioning, we used a delay paradigm consisting of a 350 ms tone (2.4 kHz, 85 dB) as a conditioned stimulus (CS) followed at its end by an electrical shock (0.5 ms, 3 x threshold, cathodal, square pulse), applied to the supraorbital nerve, as an unconditioned stimulus (US).

A total of two habituation and 10 conditioning sessions were carried out for each animal. A conditioning session consisted of 60 CS-US presentations and lasted for about 30 min. Paired CS-US presentations were separated at random by 30 ± 5 s. For habituation sessions, the CSs were presented alone, also for 60 times per session, at intervals of 30 ± 5 s (Gruart et al., 2006).

The electromyographic (EMG) activity of the orbicularis oculi muscle was recorded with Grass P511 differential amplifiers (Grass-Telefactor, West Warwick, RI, USA), at a bandwidth of 0.1 Hz-10 kHz.

#### 4.4.6. Vestibular stimulation and recording of the VOR

For vestibular stimulation, a single animal was placed on a home-made turning-table system. Its head was immobilized with the help of the implanted holding system, while the animal was allowed to walk over a running wheel. Table rotation was carried out by hand following a sinusoidal display in a computer screen. Actual rotation of the table was recorded with a potentiometer attached to its rotating axis. The animal was rotated by ± 20 deg at three selected frequencies (0.1, 0.3, and 0.6 Hz) for about ten cycles with intervals of 5 s between frequencies. The mouse right eye was illuminated with a red cold light attached to the head holding system. Eye positions during rotation were recorded with a fast infrared CCD camera (Pike F-032, Allied Technologies, Stadtroda, Germany) affixed to the turning table, at a rate of 50 or 100 pictures/s.

Recordings of head rotations and eye positions were synchronized and analyzed offline with an analog/digital converter (CED 1401 Plus, Cambridge, England) for gain and phase. Gain was computed as the averaged angular displacement of the eye (peak-to-peak) divided by the angular displacement of the head evoked experimentally. VOR gain in mice is usually << 1 (de Jeu and de Zeeuw, 2012; Stahl et al., 2006). Phase was determined as the averaged angular difference (in degrees) between peak eye position vs. peak head position (Sergaki et al., 2017; de Jeu and de Zeeuw, 2012).

#### 4.4.7. Social interaction test

To evaluate social interaction the subject (male adult mice) was allowed to move freely in a neutral cage (69 × 41 × 37cm), containing a small inverted grid box on each side. For habituation each mouse was placed in the cage for 5 min with both boxes empty to discard mice that preferred either half of the cage. In the sociability session, a strange juvenile mouse that had not previously been encountered was put underneath one of the grid inverted boxes. In the social novelty session, another stranger mouse was put inside the previously empty box. Each session lasted 10 min, and all the experiments were video recorded and analyzed by the researcher. The test estimates the time spent by the subject in close proximity to the juvenile mice, including the time during which the subject oriented its nose to the occupied box and sniffed in a distance <1 cm, and the time the subject touched the occupied box.

#### 4.4.8. Data collection and analysis of “in vivo” experiments

Recorded videos from each test were analyzed blind to the genotype and treatment. For classical eyeblink conditioning, unrectified EMG activity of the orbicularis oculi muscle, and 1-V rectangular pulses corresponding to CS and US presentations were stored digitally in a computer through an analog/digital converter (CED 1401-plus) for quantitative off-line analysis. Collected data were sampled at 10 kHz for EMG recordings, with an amplitude resolution of 12 bits. A computer program (Spike2 from CED) was used to display the EMG activity of the orbicularis oculi muscle. For experiments involving the VOR, head and eye positions were stored digitally in the same an analog/digital converter and processed off-line with the help of a MATLAB based (MathWorks, Natick, MA) home-made tracking program.

### 4.5. Statistics

The appropriate sample size was previously computed depending on the power of each test using Statgrahics Centurion XVII.2. For statistical analysis GraphPad InStat v2.05a (GraphPad Software, San Diego, CA, USA); SigmaPlot 10 (Systat Software Inc., San Jose, CA USA) and OriginPro 8.0 (OriginLab Corporation, Northampton, MA, USA) were used. For comparison between two sets of data unpaired two-tailed Student t test was used or the Welch test when the variances of the populations were significantly different. When more than two sets of data were compared, one-way ANOVA (post hoc Holm-Sidak’s method), two-way ANOVA (Bonferroni’s or Tukey’s post hoc tests) or two-way repeated measures ANOVA followed by all paired multiple comparisons procedure Holm-Sidak’s or LSD post hoc tests, were used. When variances of the populations were significantly different the Kruskal-Wallis test was used and Dunn’s as post hoc test. For details see (Gruart et al., 2006). The data are represented as the mean ± S.E.M.: \**P*<0.05, \*\**P*<0.01, \*\*\**P*<0.001. Differences were considered statistically significant when *P*<0.05 with a confidence limit of 95%.

### 4.6. Study approval

*Fmr1*KO (Strain #:003025, RRID:IMSR_JAX:003025) or WT (Strain #:000664, RRID:IMSR_JAX:000664) mice were used to establish the colonies at the Animal House Service at the Complutense University, an authorized center for the breeding of genetically modified mice. *Fmr1*KO and mice and WT littermate were used in this study. These mice were also supplied to the Pablo de Olavide Animal House (Sevilla, Spain). Experiments were carried out in accordance with guidelines of the European Union Council (2010/276:33-79/EU) and Spanish (BOE 34:11370-421, 2013) regulations for the use of laboratory animals in chronic studies and, in addition, were approved by the Ethics Committee of Comunidad de Madrid (PROEX 012/18) and of the Junta de Andalucía (code 06/04/2020/049).

## AUTORSHIP CONRIBUTION

J.S.-P., M.T., N.G.-F., R.M., A.G., J.M.D.-G. conceived and designed the experiments; R.M., N.G.-F., A.S-P., M.J.O.-G., M.L.L.-L., J.C.L.-R., A.G. and J.M.D.-G. acquired and analyzed the data; J.S.-P., J.M.D.-G. wrote the paper.

## DATA AVAILABILITY

All data generated or analysed during this study are included in the manuscript and Supplementary Material. Source data files for main figures and supplementary figures have also been provided.

## ACKNOWLEDGEMENTS

We thank ML. García and M. González from the electron microscopy facility at the Universidad Complutense de Madrid for technical support, and J.M. Gonzalez Martin and M. Sanchez Enciso from Universidad Pablo de Olavide for their help in the performed experiments. Miguel Delicado Miralles, a visiting fellow from the INA-CSIC of Alicante, also helped in some in vivo experiments. We also thank Dr Bartolomé-Martín (Departamento de Ciencias Médicas Básicas (Fisiología), Universidad de la Laguna) for his help with statistics. We thank Mark Sefton for editorial assistance and Dr I Galve-Roperh (Facultad de Biología at UCM) for facilitating the chamber to perform the skilled reaching tests.

## SUPPLEMENTARY MATERIAL

### SUPPLEMENTARY FIGURE LEGENDS

**Figure S1.**
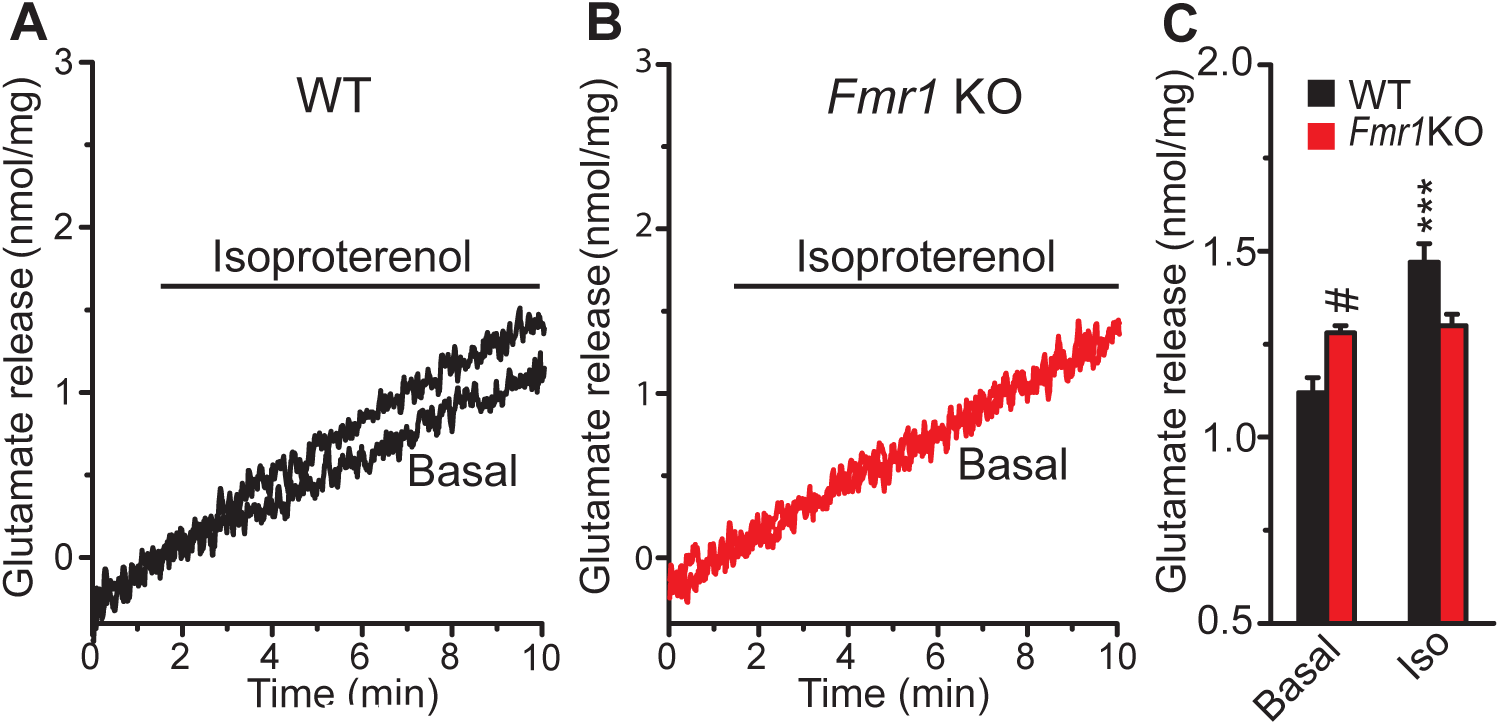
Lack of isoproterenol-induced potentiation of glutamate release in *Fmr1* KO cerebellar synaptosomes. (A, B) Mean traces from WT (A) and *Fmr1* KO (B) cerebellar synaptosomes showing spontaneous release of glutamate in the presence of the Na^+^ channel blocker tetrodotoxin (1 μM, TTx), and in the presence or absence (control) of isoproterenol (100 μM). (C) Diagram summarizing the isoproterenol effect on glutamate release in the aforementioned conditions from WT (n=8/3 synaptosomal preparations: *P*=0.0005) and *Fmr1* KO synaptosomes (n=7/3 preparations: *P*= 0.719). Basal release from *Fmr1* KO vs WT synaptosomes (^#^*P*=0.011). The data represent the mean ± SEM. Unpaired Student’s *t* test in C. \*\*\**P*<0.001

**Figure S2.**
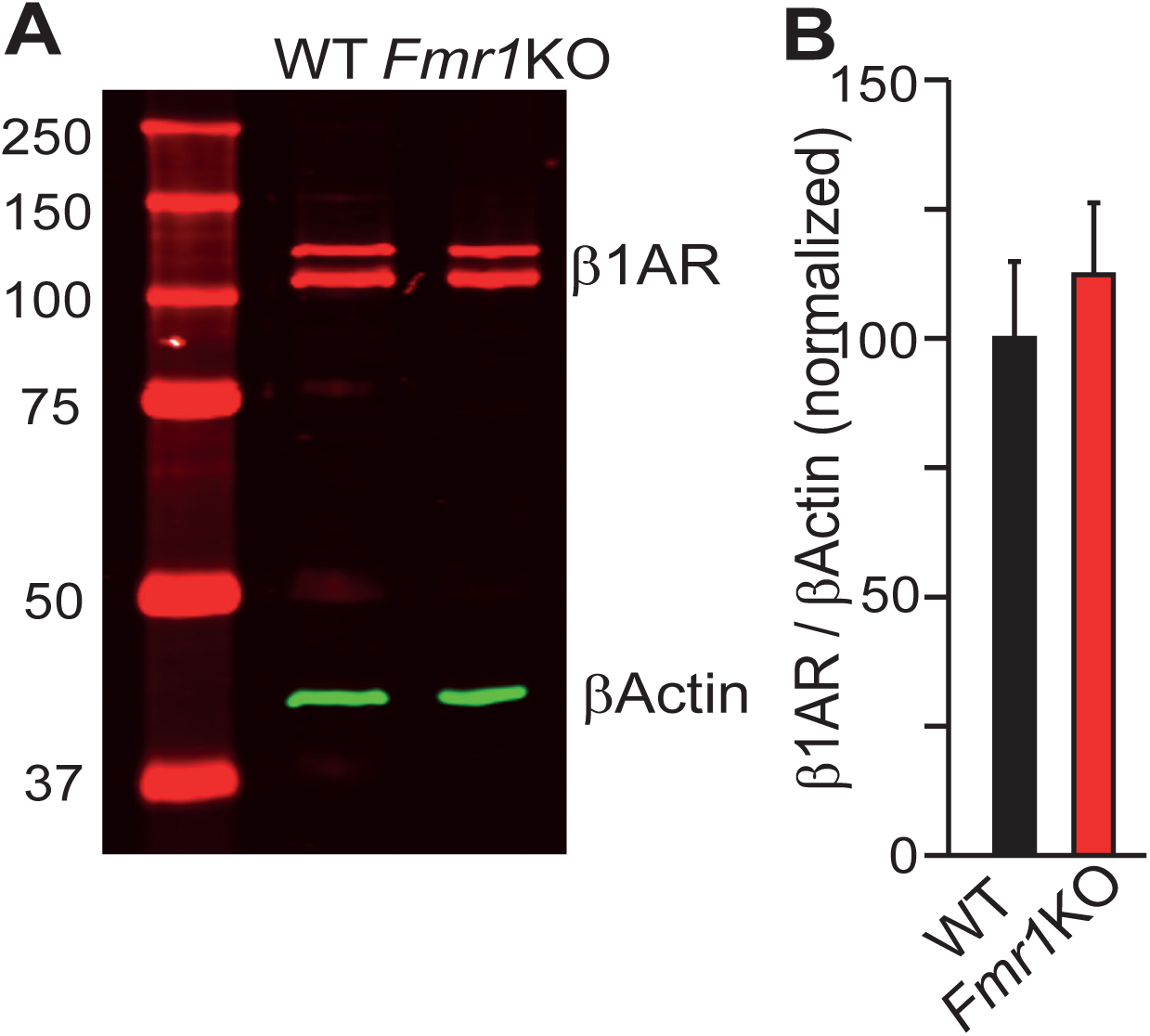
The expression of β-AR is not altered in *Fmr1* KO cerebellar synaptosomes. (A) Western blot analysis of β1-AR in the P_2_ crude synaptosomes fraction from WT and *Fmr1* KO mice. (B) The data were normalized to the WT values (n=3/3preparations (*Fmr1* KO, n=3/3 preparations: *P*=0.5773). The data represent the mean ± SEM. Unpaired Student’s t test in B.

**Figure S3.**
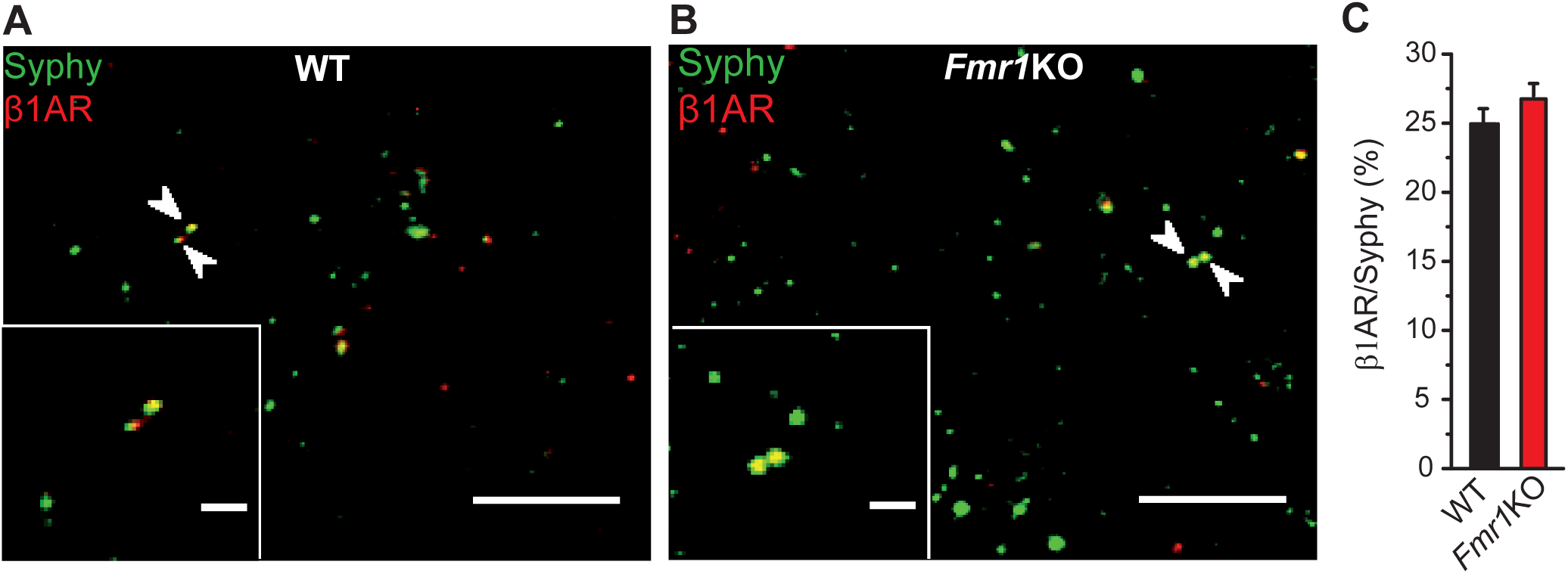
The distribution of β-AR is not altered in *Fmr1* KO cerebellar synaptosomes. (A,B) Quantification of β-AR expressing cerebellar synaptosomes. Immunofluorescence of WT (A) and *Fmr1* KO (B) synaptosomes stained with antibodies against β_1_-AR and the vesicular marker synaptophysin. (C) Co-expression of β-AR/synaptophysin in WT (24.9 ±1.1%, n=16,804/63 fields/2 preparations) and in *Fmr1* KO synaptosomes (26.7 ±0.9%, n=18,712/75 fields/2 preparations, *P*=0.200). Scale bar in A and B, 5 μm. The data represent the mean ± SEM. Unpaired Student’s t test.

**Figure S4.**
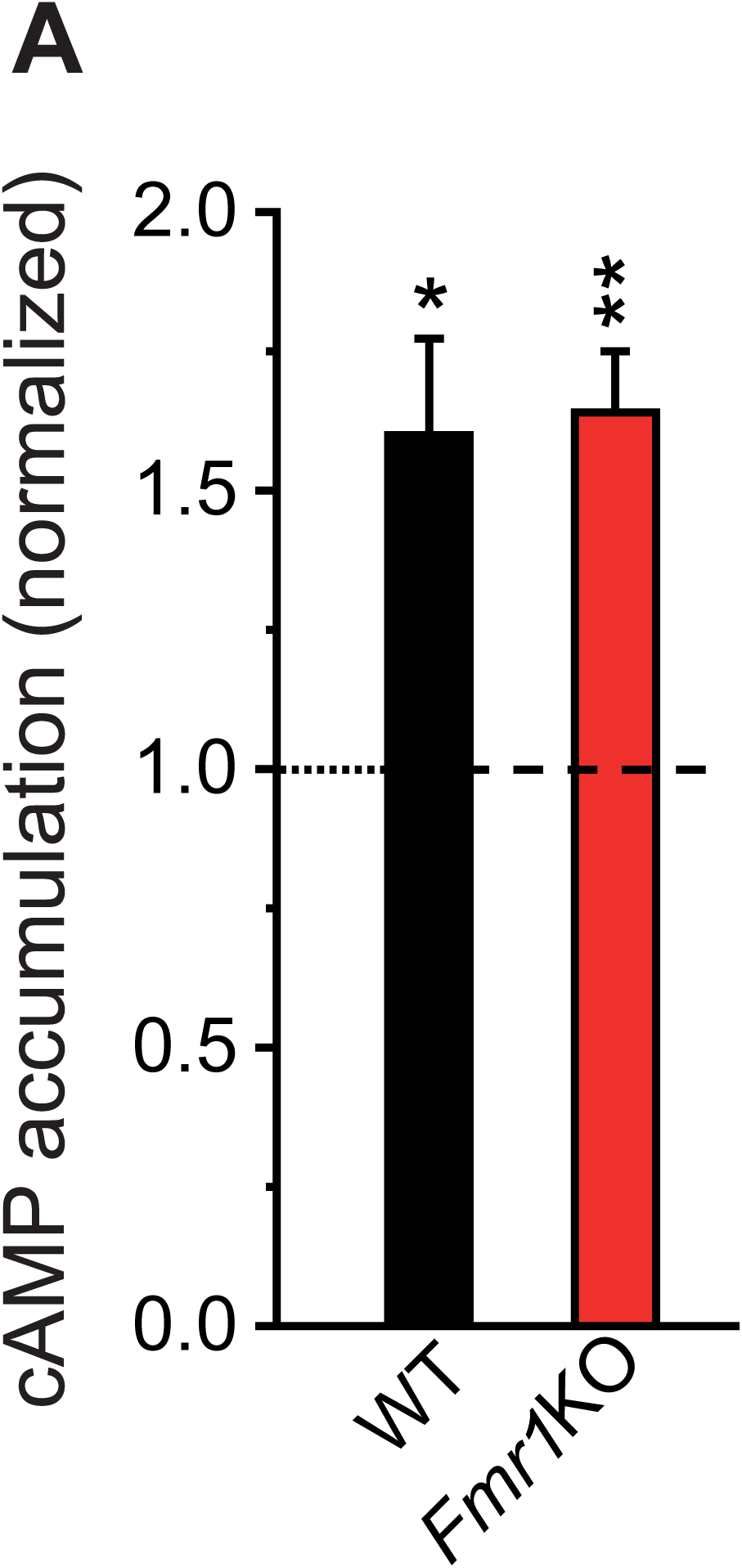
The isoproterenol induced generation of cAMP is not altered in *Fmr1* KO cerebellar synaptosomes. The effect of isoproterenol on the cAMP levels in WT (n=14/3 preparations: *P*=0.032) is similar (*P*=0.848) to that in *Fmr1* KO synaptosomes (n=15/3 preparations: \*\**P*=0.0043, unpaired Student’s t test). The data represent the mean ± SEM. \**P*<0.05. \*\**P*<0.01.

**Figure S5.**
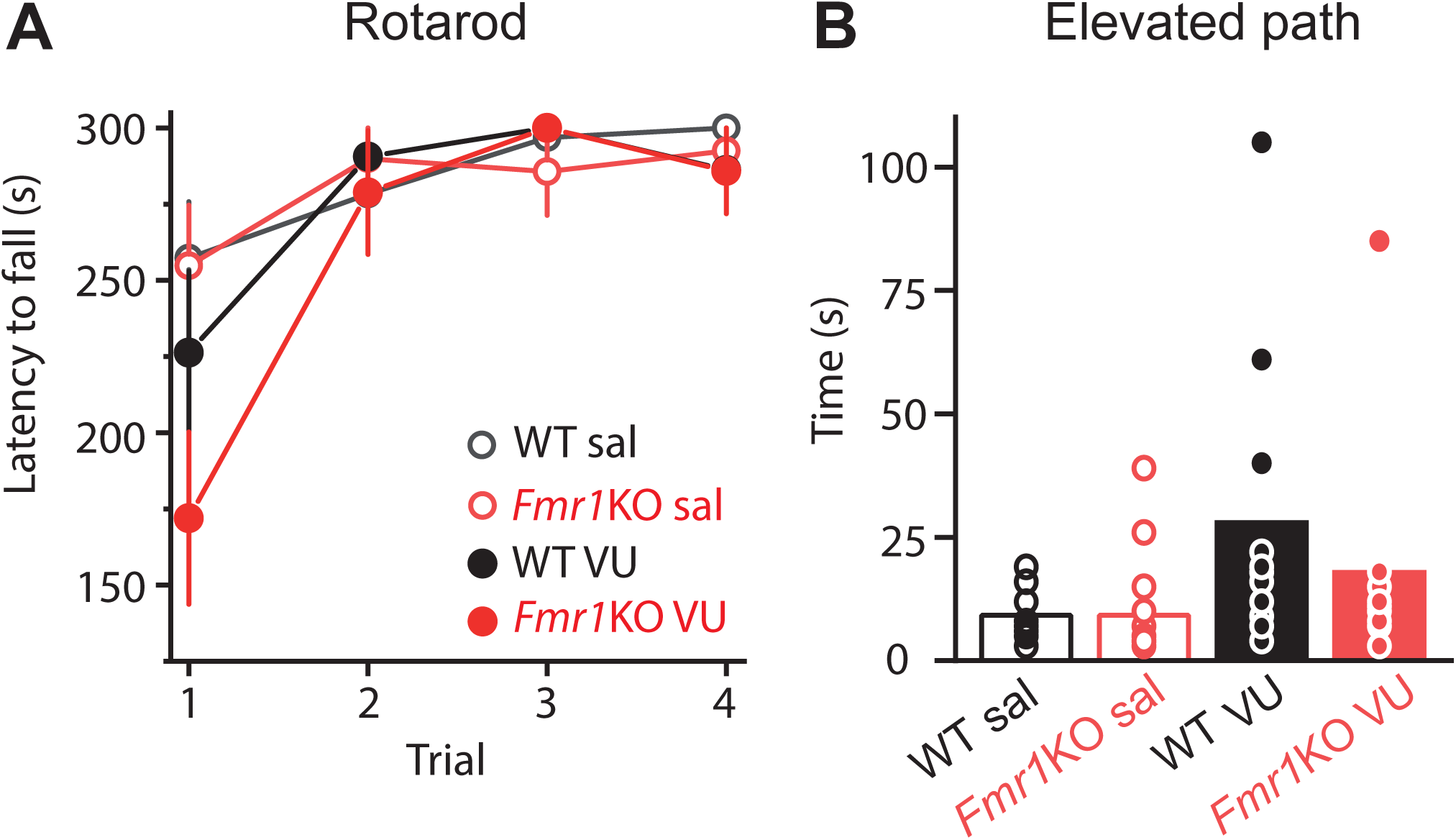
*Fmr1* KO performance in rotarod and elevated path is unchanged. (A), Latency to fall of the mouse in an accelerating rotarod. WT sal (n=11), *Fmr1*KO sal (n=14) *P*>0.05, WT VU (n=10) *P*>0.05, and *Fmr1*KO VU (n=11) *P*>0.05. Kruskal-Wallis followed by Dunn’s test. All comparisons were to WT sal. (B), In the elevated path the time spent to walk from the center of a 5 cm wide bar to one of its ends is measured. WT sal, *Fmr1*KO sal *P*>0.05, WT VU *P*>0.05, and *Fmr1*KO VU *P*>0.05, Kruskal-Wallis followed by Dunn’s test. All comparisons were to WT sal. The data represent the mean ± SEM.

